# Jak-Stat pathway induces Drosophila follicle elongation by a gradient of apical contractility

**DOI:** 10.1101/205187

**Authors:** Hervé Alégot, Pierre Pouchin, Olivier Bardot, Vincent Mirouse

**Affiliations:** GReD Laboratory, Université Clermont Auvergne, CNRS, INSERM, F-63000 Clermont-6 Ferrand, FRANCE

## Abstract

Tissue elongation and its control by spatiotemporal signals is a major developmental question. Currently, it is thought that *Drosophila* ovarian follicular epithelium elongation requires the planar polarization of the basal domain cytoskeleton and of the extra-cellular matrix, associated with a dynamic process of rotation around the anteroposterior axis. Here we show, by careful kinetic analysis of *fat2* mutants, that neither basal planar polarization nor rotation is required during a first phase of follicle elongation. Conversely, a JAK-STAT signaling gradient from each follicle pole orients early elongation. JAK-STAT controls apical pulsatile contractions, and Myosin II activity inhibition affects both pulses and early elongation. Early elongation is associated with apical constriction at the poles and oriented cell rearrangements, but without any visible planar cell polarization of the apical domain. Thus, a morphogen gradient can trigger tissue elongation via a control of cell pulsing and without planar cell polarity requirement.

**Impact Statement:** Follicle elongation does not rely solely on the basal side of the cells but also requires a mechanism integrating a developmental cue with a morphogenetic process involving their apical domain.

## Introduction

Tissue elongation is an essential morphogenetic process that occurs during the development of almost any organ. Therefore, uncovering the underlying molecular, cellular and tissue mechanisms is an important challenge. Schematically, tissue elongation relies on at least three determinants. First, the elongation axis must be defined by a directional cue that usually leads to the planar cell polarization (pcp) of the elongating tissue. Second, a force producing machinery must drive the elongation and this force can be generated intrinsically by the cells within the elongating tissue and/or extrinsically by the surrounding tissues. Finally, such force induces tissue elongation via different cellular behaviors, such as cell intercalation, cell shape modification, cell migration or oriented cell division. This is exemplified by germband extension in *Drosophila* embryo where Toll receptors induce Myosin II planar polarization, which drives cell rearrangements (Bertet et al 2004; Irvine, & Wieschaus 1994; Blankenship et al 2006; Paré et al 2014).

In the last years, *Drosophila* egg chamber development has emerged as a powerful model to study tissue elongation (Bilder, & Haigo 2012; Cetera, & Horne-Badovinac 2015). Each egg chamber (or follicle) consists of a germline cyst that includes the oocyte, surrounded by the follicular epithelium (FE), a monolayer of somatic cells. The FE apical domain faces the germ cells, while the basal domain is in contact with the basement membrane. Initially, a follicle is a small sphere that progressively elongates along the anterior-posterior (AP) axis, which becomes 2.5 times longer than the mediolateral axis (aspect ratio (AR) = 2.5), prefiguring the shape of the fly embryo.

All the available data indicate that follicle elongation relies on the FE. Specifically, along the FE basal domain F-actin filaments and microtubules become oriented perpendicularly to the follicle AP axis (Gutzeit 1990; Viktorinová, & Dahmann 2013). The cytoskeleton planar polarization depends on the atypical cadherin Fat2 via an unknown mechanism (Viktorinová et al 2009; Viktorinová, & Dahmann 2013; Chen et al 2016). Fat2 is also required for a dynamic process of collective cell migration of all the follicle cells around the AP axis until stage 8 of follicle development. This rotation reinforces F-actin planar polarization and triggers the polarized deposition of extracellular matrix (ECM) fibrils perpendicular to the AP axis (Haigo, & Bilder 2011; Lerner et al 2013; Viktorinová, & Dahmann 2013; Cetera et al 2014; Isabella, & Horne-Badovinac 2016; Aurich, & Dahmann 2016). These fibrils have been proposed to act as a molecular corset, mechanically constraining follicle growth along the AP axis during follicle development (Haigo, & Bilder 2011). Additionally, Fat2 is required for the establishment of a gradient of basement membrane (BM) stiffness at both poles at stage 7-8 (Crest et al 2017). This gradient also depends on the morphogen-like activity of the JAK-STAT pathway and softer BM near the poles would allow anisotropic tissue expansion along the A-P axis (Crest et al 2017). After the end of follicle rotation, F-actin remains polarized in the AP plane during stage 9 to 11 and follicular cells (FCs) undergo oriented basal oscillations that are generated by the contractile activity of stress fibers attached to the basement membrane ECM via integrins ((Bateman et al 2001; Delon, & Brown 2009; He et al 2010).

Nonetheless, in agreement with recently published observations, we noticed that a first phase of follicle elongation does not require *fat*2 and the planar polarization of the basal domain (Aurich, & Dahmann 2016). We therefore focused on this phase, addressing the main three questions which are how the follicle elongation axis is defined, what is the molecular motor triggering elongation in a specific axis, and how FCs behave during this phase.

## Results

### Polar cells define the axis of early elongation

We analyzed the follicle elongation kinetics in *fat2^58D^* mutants, which block rotation and show a strong round egg phenotype. Follicle elongation is normal in *fat2* mutants during the first stages (3 to 7) with an AR of 1.6 (Fig 1a-d). Thus, at least two mechanistically distinct elongation phases control follicle elongation, a first phase (stage 3 to 7) independent of *fat2*, rotation and ECM basal polarization, and a later one (stage 8 to 14) that requires *fat2*. This observation is consistent with the absence of elongation defect of clonal loss-of-function of *vkg* before stage 7-8 (Bilder and Haigo, 2011).

**Figure 1:**
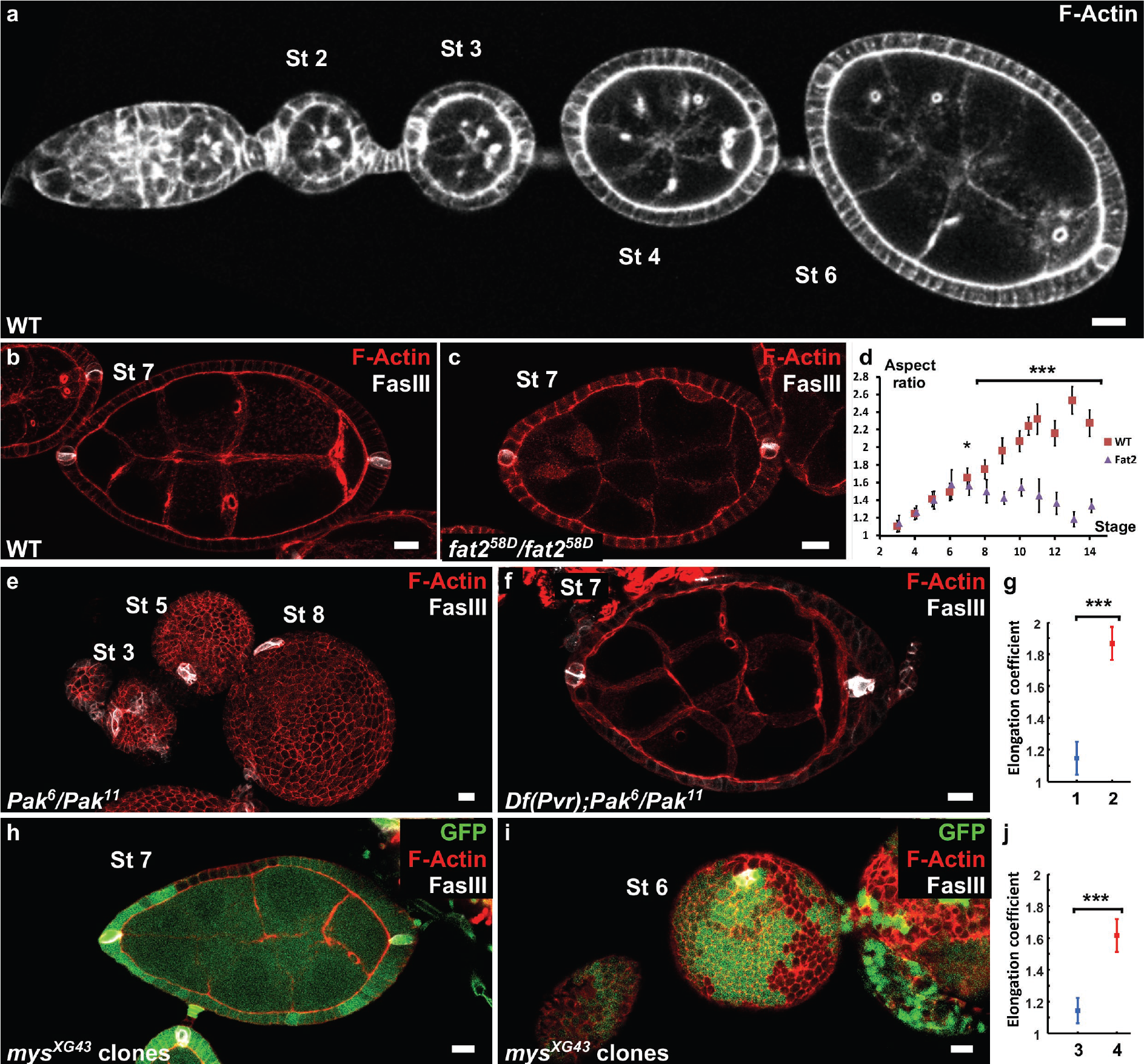
Polar cells determine the axis of early elongation. a) WT ovariole illustrating follicle elongation during the early stages of oogenesis (stage 2 to 6). b) Optical cross-section of a stage 7 WT follicle stained with Faslll, a polar cell marker (white), and F-actin (red). c)Stage 7 *fat2* mutant follicle stained with FasIII (white) and DE-Cad (red). d) Elongation kinetics of WT and fat2 mutant follicles. e) Z-projection of a *Pak* mutant ovariole. Round follicles have only one cluster of polar cells (stage 5 and 8 follicles) or two non-diametrically opposed clusters (stage 3 follicle)._f) Removing a copy of *Pvr* restores early elongation and polar cell position in *Pak* mutants. g) elongation coefficient of *Pak*^6^/*Pak*^11^,*Df*(*Pvr*)/+ follicles, affecting (1) or not (2) polar cell positioning. h,i) view of a *mys* mutant clone (GFP-negative) in a mosaic follicle shoing h) normal polar cell positioning and no elongation defect and i) abnormal polar cell positioning and early elongation defect. j) elongation coefficient of follicles containing mutant clones for *mys* affecting (3) or not (4) polar cell positioning. (p *< 0.05,**<0.01,***<0.001). On all picture scale bar is 10 μm.

To try to identify the mechanism regulating the early phase of follicle elongation, we first analyzed trans-heterozygous *Pak* mutant follicles, Which never elongate (Conder et al 2007) (fig. 1e). The *Pak* gene encodes a Pak family serine/threonine kinase that localizes at the FE basal domain. *Pak* mutants also show many other abnormalities, such as the presence of more than one germline cyst and abnormal interfollicular filaments ((Vlachos et al 2015) and not shown). Interfollicular cells derive from prepolar cells that also give rise to the polar cells, which prompted us to analyze distribution of the latter using the specific marker FasIII (Bastock, & St Johnston 2008; Horne-Badovinac, & Bilder 2005). Polar cells are pairs of cells that differentiate very early and are initially required for germline cyst encapsulation (Grammont, & Irvine 2001). They also play the role of an organizing center for the differentiation of FC sub-populations during midoogenesis (Xi et al 2003). In WT follicles, polar cells are localized at the follicle AP axis extremities (Fig 1b). Conversely, in *Pak* mutants, we observed a single polar cell cluster or two clusters close to each other (Fig 1e). This suggests that *Pak* is required for polar cell positioning, though a role in their specification or survival cannot be excluded, which in turn could play a role in defining the elongation axis. Some dominant suppressors of the *Pak* elongation defect have been identified, including PDGF- and VEGF-receptor related (Pvr), although the reason for this suppression is unknown (Vlachos, & Harden 2011). By using flies heterozygous for a *Pvr* allele and mutant for *Pak*, we observed that normal positioning of polar cells is frequently but not always restored (Fig 1f and 1S1c). We quantitatively compared the elongation of those two situations, normal or abnormal polar cells, by plotting the long axis as a function of the short axis for previtellogenic stages (before stage 8) and determined the corresponding regression line (Fig 1S1d). We defined an elongation coefficient that corresponds to the slope of this line and for which a value of 1 means no elongation. This method allows to quantify elongation independently of any bias that could be introduced by stage determination approximation due to aberrant follicle shape or differentiation. Moreover, focusing on previtellogenic stages allows excluding genotypes that affect only the late elongation phase. It is exemplified with *fat2* mutant that does not induce significant defects if we include only stage 3 to 7 follicles (previtellogenic), but does show a difference if we include stage 8 (Fig 1S1a,b). The statistical comparison of the elongation coefficients clearly shows that restoring polar cell position by removing one copy of *Pvr* in *Pak* mutants strongly rescues follicle elongation (Fig 1g and 1S1c,d).

Although it has not been fully demonstrated in this context, Pak often works as part of the integrin signaling network and mosaic follicles containing FC clones mutant for *myospheroid* (*mys*), which encodes the main fly (β-integrin subunit, also show a round follicle phenotype at early stages (Haigo, & Bilder 2011). We noticed that in some follicles containing *mys* mutant clones, polar cells are mispositioned, a defect generally observed when at least one polar cell is mutant. As in *Pak* mutants, the two polar cell clusters are not diametrically opposed (Fig 1S1e), or a single cluster is observed (Fig 1i, movie S1). Importantly, the polar cell positioning defect is associated with the round follicle phenotype (Fig 1j, and 1S1f). Conversely, in mosaic follicles in which polar cell positioning was not affected, the round egg phenotype is never observed at early stages, even with large mutant clones (Fig 1h and 1S1f, movie S2). In agreement, the elongation coefficient of the latter cases is much higher than the ones with abnormal polar cells (Fig 1j). Thus, together, these results indicate that *pak* and *mys* mutants are not required for the early phase of elongation once polar cells are well-placed and thus affect this phase indirectly. They also strongly suggest that polar cells are required to define the follicle elongation axis.

### A JAK-STAT gradient from the poles is the cue for early elongation

Once the follicle is formed, polar cells are important for the differentiation of the surrounding FCs. From stage 9 of oogenesis, FCs change their morphology upon activation by Unpaired (Upd), a ligand for the JAK-STAT pathway, exclusively produced by polar cells throughout oogenesis (Silver, & Montell 2001; Xi et al 2003; McGregor et al 2002). To identify the FCs in which the JAK-STAT pathway is active, we used a reporter construct in which GFP transgene expression is controlled by STAT binding repeat elements in the promoter (Bach et al 2007). During the early stages of oogenesis, the pathway is active in all the main body FCs (Fig 2b). Moreover, we observed differences in GFP expression level (and thus STAT activity) between the poles and the mediolateral region, starting at about stage 3, concomitantly with the beginning of elongation (Fig 2b,h). At later stages (5 to 7), it leads to the formation of a gradient of STAT activity, as indicated by the strong GFP expression at each pole and the very weak or no signal in the large mediolateral part of each follicle (Fig 2b,h, and 2S1a). Thus, the spatiotemporal pattern of JAK-STAT activation is consistent with a potential role of this pathway in follicle elongation.

**Figure 2:**
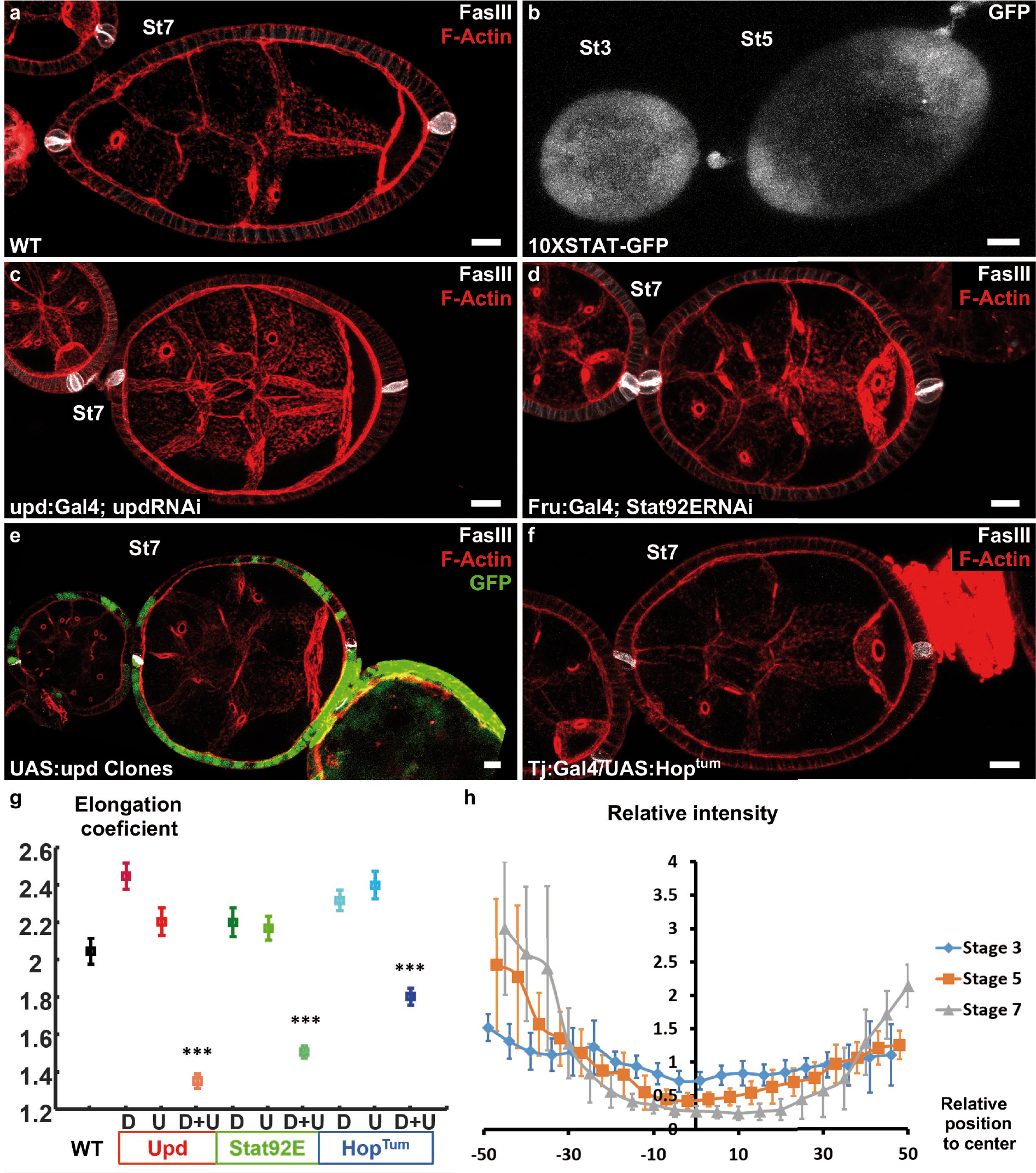
Upd is a polarizing cue for early elongation. a) Optical cross-section of a WT stage 7 follicle stained with FasIII (polar cell marker, white) and F-actin (red). b) Expression of the 10xStatGFP reporter shoing the progressive formation of a STAT gradient at each pole. Early elongation is affected by knocking don c) *upd* in polar cells or d) *Stat92E* in the anterior and posterior follicular cells.Early elongation is also affected by e) clonal ectopic expression of upd (GFP-positive cells) and by f) expression of a Hop gain of function mutant in all follicular cells. g) Quantification of the elongation coefficient in WT and the different JAK-STAT mutants (loss and gain of function) during early and intermediate stages of elongation (D, Driver; U, UAS line) (p *< 0.05,**<0.01,**<0.001). h) Quantification of the Stat activity gradient at stage 3,5 and 7 using the 10XSTATGFP reporter. A gradient is already visible at stage 3 and get more visible until stage 7. On all picture scale bar is 10 μm. Relative intensity = intensity at a given position/mean intensity of measured signal.

The key role of JAK-STAT signaling during follicle formation precluded the analysis of elongation defects in large null mutant clones (McGregor et al 2002). Therefore, we knocked-down by RNAi the ligand *upd* and the most downstream element of the cascade, the transcription factor *Stat92E*, both efficiently decreasing the activity of the pathway in the follicular epithelium (Fig 2a,c,d,g and 2S1). *Upd* knock-down was performed either using upd:Gal4 that is specifically expressed in the polar cells (Khammari et al 2011), or tj:Gal4 expressed in all FCs, and then analyzed only follicles that contained one germline cyst and correctly placed polar cells. At early stages, with both drivers, such follicles are significantly rounder than control follicles (Fig 2a,c,g and 2S1e). It indicates a role for JAK-STAT pathway in early elongation and confirms the causal link between polar cells and early elongation. Moreover, knock-down of *Stat92E* using a driver specifically expressed at the poles (Fru:Gal4) also affects early elongation (Fig 2d,g, and 2S1e), suggesting a transcriptional control of elongation by JAK-STAT (Borensztejn et al 2013). These results are the first examples of loss of function with an effect only on early elongation independent of polar cells position, and indicate that Upd secreted by the polar cells and JAK-STAT activation in FCs are both required for follicle elongation. Moreover, clonal ectopic *upd* overexpression completely blocks follicle elongation, without affecting polar cell positioning (Fig 2e), demonstrating that Upd is not only a prerequisite for the elongation but the signal that defines its axis (n = 20). Similarly, general expression of Hop^Tum^, a gain of function mutation of fly JAK, disrupting the pattern of JAK-SAT activation, also affects follicle elongation (Fig 2f,g, and 2S1e). Thus, the spatial control of the JAK-STAT pathway activation is required for follicle elongation. Altogether, these results show that Upd secretion by polar cells and the subsequent gradient of JAK-STAT activation act as a developmental cue to define the follicle elongation axis during the early stages of oogenesis.

### MyosinII activity drives apical pulses and early elongation

Once the signal for elongation identified, we aimed to determine the molecular motor driving this elongation, which in many morphogenetic contexts is MyosinII (MyoII)(Heisenberg, & Bellaïche 2013; Lecuit et al 2011). The knock-down in all FCs of *spaghetti squash* (*sqh*), the MyoII regulatory subunit, leads to a significant decrease in the elongation coefficient and follicle AR from stage 4 (Fig 3a-b and 3S1b c) indicating that MyoII is the motor of early elongation. We have shown that the rotation and the planar polarization of the basal actomyosin is not involved in early elongation. Moreover, at these stages, MyoII is strongly enriched at the apical cortex suggesting that its main activity is on this domain of the FCs (Fig 3S1a and 5c) (Wang, & Riechmann 2007). We therefore looked at MyoII on living follicles focusing on the apical side and found that it is highly dynamic (movie S3). In *Drosophila*, transitory medio-apical recruitment of actomyosin usually drives apical pulses (Martin et al 2009, Martin, & Goldstein 2014). Accordingly, using a GFP trap line for Bazooka (Baz-GFP), which concentrates at the *zonula adherens* and marks the periphery of the apical domain, we observed that the transient accumulation of MyoII is associated with a contraction of this domain, followed by a relaxation when MyoII signal decreases (Fig 3c-e, Movie S4). Although we did not find a clear period because cells can pause for a variable time between two contractions, the approximate duration of a pulse was about three minutes.

**Figure 3:**
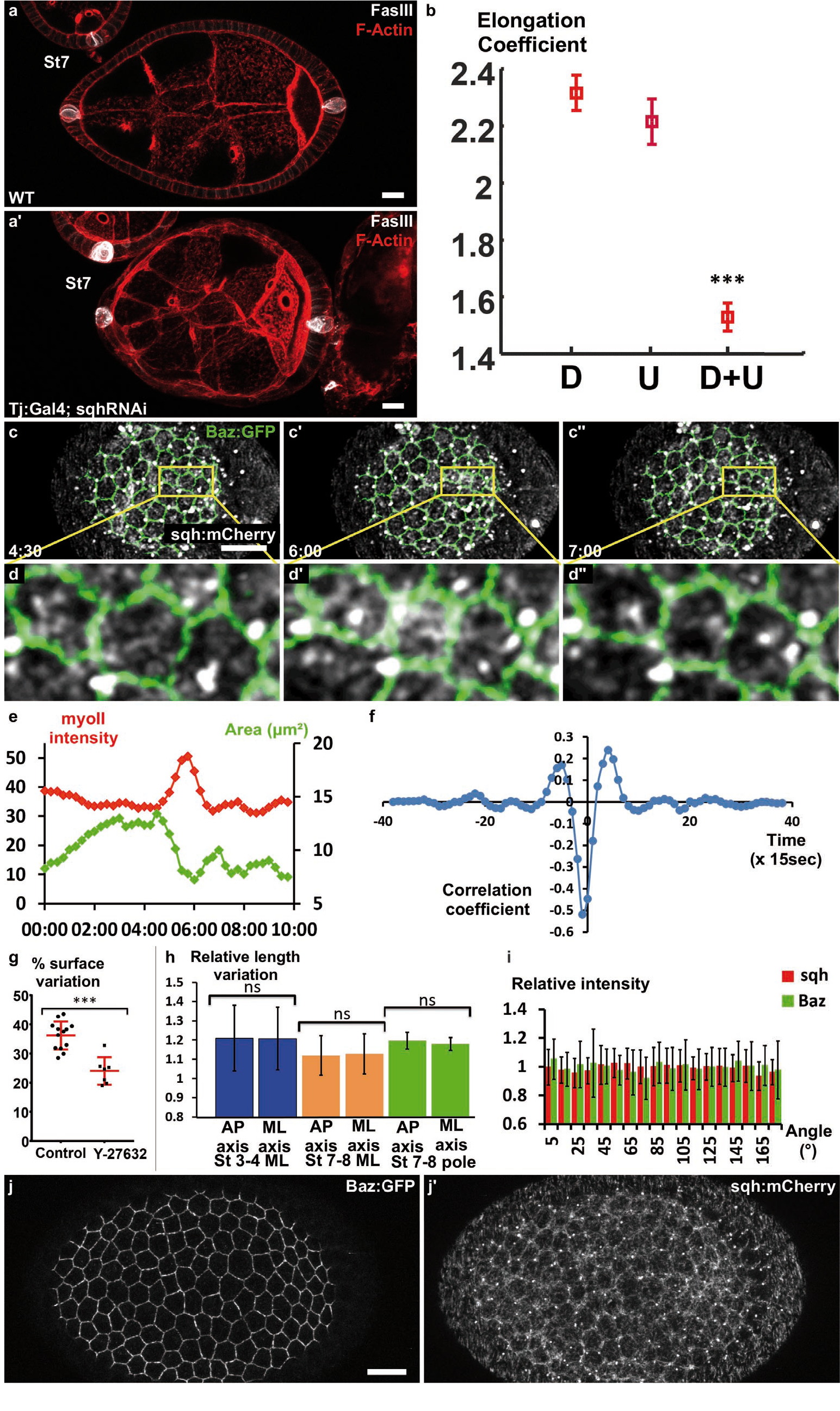
Myosin II is required for early elongation and apical pulses. a) WT and a’) *sqh* knock-down stage 7 follicles stained for F-actin (red) and FasIII (White). b) elongation coefficient of WT or *sqh* knock-down follicles during the early elongation phase (D, Driver; U, UAS line). c) Fluorescence video-microscopy images of a stage 4 WT follicle that expresses BAZ-GFP and Sqh-mCherry. d) Higher magnification of the area highlighted in (c) shoing a pulsing cell. e) Quantification of the cell apical surface (green) in the cell shown in (d) and of Sqh signal intensity in the apical area (red) over E Dh time. f) Cross correlation analysis over time of apical surface and Sqh apical signal intensity based on 86 cells from 6 follicles at stage 3-4. g) Incubation with the Rok inhibitor Y-27632 strongly reduces pulse activity in stages 3 to 5 WT follicles. Red bars represent mean and +/− sd, (n≥7 follicles) h) Quantification of the length variation of the follicle cell AP and mediolateral axes during pulses indicates that pulses are isotropic. i) Quantification of the relative BAZ-GFP and Sqh-mCherry signal intensity in cell bounds in function of their angle relative to the AP axis (n=44 follicles). Relative intensity is given over the mean bond intensity. j) Fixed stage 7 WT follicle that expresses GFP-Baz and Sqh-mCherry. On all picture scale bar is 10 μm. (p ***<0.001)

Cross correlation analysis on many cells from several follicles (n=86) confirms the association between MyoII and pulses and reveals that Sqh accumulation slightly precedes the reduction of the apical surface, arguing that it is the motor responsible for these contractions (Fig 3f). Inhibiting Rho kinase (rok) activity, the main regulator of MyoII, using Y-27632, reduces follicle cells surface variation by ~30% (Fig 3g). Thus, MyoII drives apical pulsing during early stages. Consequently, we asked whether and how apical pulses could induce elongation. From stage 9, basal pulses, which are important for the second phase of elongation, have been shown to be anisotropic (He et al 2010). However, quantification of axis length variations showed that the apical pulses were isotropic, both in the mediolateral and polar regions (Fig 3h). Tissue elongation is often associated with tissue planar cell polarization, we therefore investigated whether Myosin II and Baz showed exclusive cortical planar polarization, as demonstrated for instance during germband extension (Bertet et al 2004; Zallen, & Wieschaus 2004). Consistent with the isotropic nature of the pulses, we failed to detect any oriented enrichment of these proteins, indicating the absence of noticeable apical planar cell polarization of the motor generating early elongation (Fig 3i,j). Altogether, these data indicate that MyoII induces apical pulses and early elongation. Nonetheless, neither the isotropic nature of the pulses nor MyoII localization explains how the pulses could induce elongation.

### JAK-STAT induces a gradient of apical pulses

Our previous results suggest that pulses do not provide an explanation for elongation at a local cellular scale and we therefore analyzed their spatiotemporal distribution at the tissue scale to determine whether they present a specific tissue pattern. Based on the JAK-STAT activity gradient we hypothesized that cells in the mediolateral part of the follicles should progressively change their behavior during follicle growth. We therefore monitored the mediolateral region of stage 3 and 7 follicles. At stage 3, cells undergo contractions and relaxations asynchronously (Fig 4a, Movie S5). At stage 7, cells were much less active (Fig 4c, Movie S6). This difference was confirmed by monitoring the variation of the relative apical surface of individual cells (Fig 4e) or a whole population (Fig 4f) (40% of mean variation at stage 3 and only about 15% in the equatorial part at stage 7). Quantification of the average variation of the apical cell surface in a series of follicles indicates that the pulsing amplitude gradually decreases in mediolateral from stage 3 to stage 8 (Fig 4S1a). This correlation between JAK-STAT activity and pulsing activity in the mediolateral region prompted us to develop a method to visualize the poles of living follicles, which has never been done before (see methods). We managed to image the poles of stage 3 to 4 and stage 7 to 8 follicles and in both cases the pulse activity is high (Fig 4b,d-f, Movie S7, S8). Finally, the analysis of slightly tilted stage 7 to 8 follicles clearly revealed a gradient of pulse intensity emanating from the pole (Fig 4g and 4S1b). Thus, pulse intensity distribution is similar in space and time to the JAK-STAT activity gradient. Moreover, the cell pulse amplitude is significantly reduced in the mediolateral region of stage 3-4 and near the poles of stage 7-8 *upd* RNAi follicles (Fig 4h,i, Movie S9, S10), indicating that JAK-STAT regulates FC apical pulsatory activity. Finally, we found that clonal ectopic activation of JAK is sufficient to increase pulse intensity in the mediolateral region of stage 7 to 8 follicles compared to similar control clones (Fig 4j, movies S11 and S12). Together, these results show that JAK-STAT pathway has an instructive role in controlling the intensity of FC apical pulses, leading to a specific spatiotemporal pattern breaking follicle symmetry in each hemisphere.

**Figure 4:**
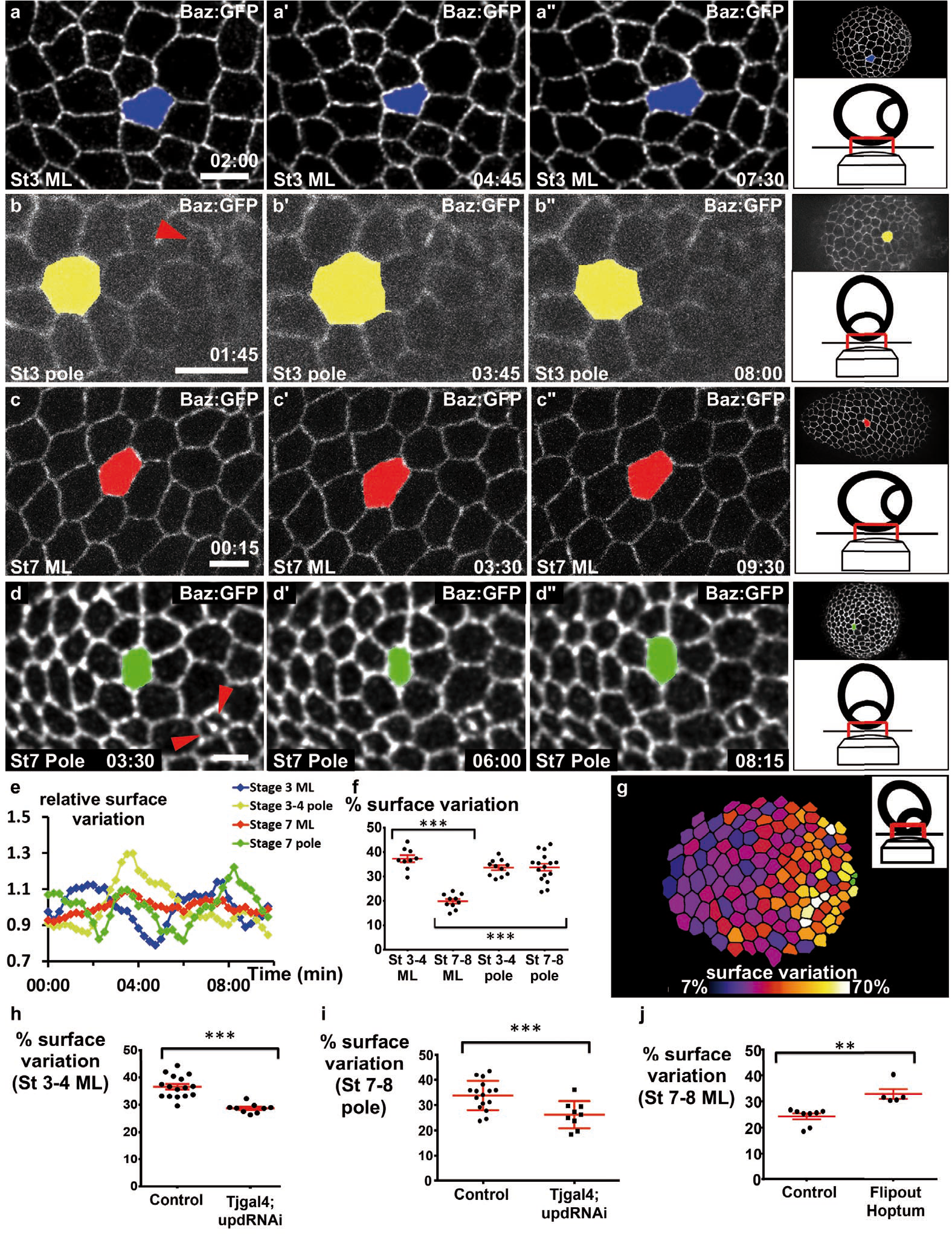
JAK-STAT induces a double gradient of pulses. a) to d) images from movies of the mediolateral region of (a) stage 3 and (c) stage 7 BAZ-GFP expressing follicles, or of the area near the polar cells (red arroheads) of (b) stage 3 and (d) stage 7 follicles. Scale bars:10 pm e) Surface variation of individual cells (examples shown in a to d) in function of time (ML: mediolateral). The surface of each cell is divided by its average surface over time. f) Mean percentage of apical surface variation depending on stage and position (n≥9 follicles). g) Colour-coding of pulse intensity of a representative stage 7 follicle (tilted view from the pole, see schematic image in insert) reveals an intensity gradient from the polar cells (in green) to the mediolateral region. h) to j) Mean percentage of apical surface variation in the mediolateral region of (h) stage 3 to 5 follicles and (j) stage 7 to 8 follicles and (i) at the pole of stage 7 to 8 follicles for the indicated genotypes. h and i) n≥9, j) n≥5 (p *<0.01,***<0.001,Red bars represent mean and +/− sd)

### Myosin II is required at the poles but is not controlled by JAK-STAT

Since JAK-STAT pathway and MyoII are both important for apical pulses, we studied their functional relationship. The apical level of the Myosin II active form, visualized by its phosphorylation, is significantly reduced by 18% in *STAT92E* null mutant clones on young follicles, compared to WT surrounding cells (n=17 clones, p<0,001), which may suggest that MyoII activity is regulated by JAK-STAT signaling (Fig 5a). However, clonal gain of function of JAK in the region where the JAK-STAT pathway is normally inactive (mediolateral at stage 7-8) does not increase the apical phosphorylation level of MyoII (Fig 5b). Moreover, analysis of the global pattern of apical MyoII phosphorylation does not reveal any gradient between poles and mediolateral regions (Fig 5c,d). Altogether these data indicate that MyoII activation by phosphorylation is independent of JAK-STAT signaling and that JAK-STAT regulates pulses by another means, which might be required for efficient apical recruitment of MyoII. Thus, although JAK-STAT and Myosin II are both required for early elongation, they control pulses in parallel.

**Figure 5:**
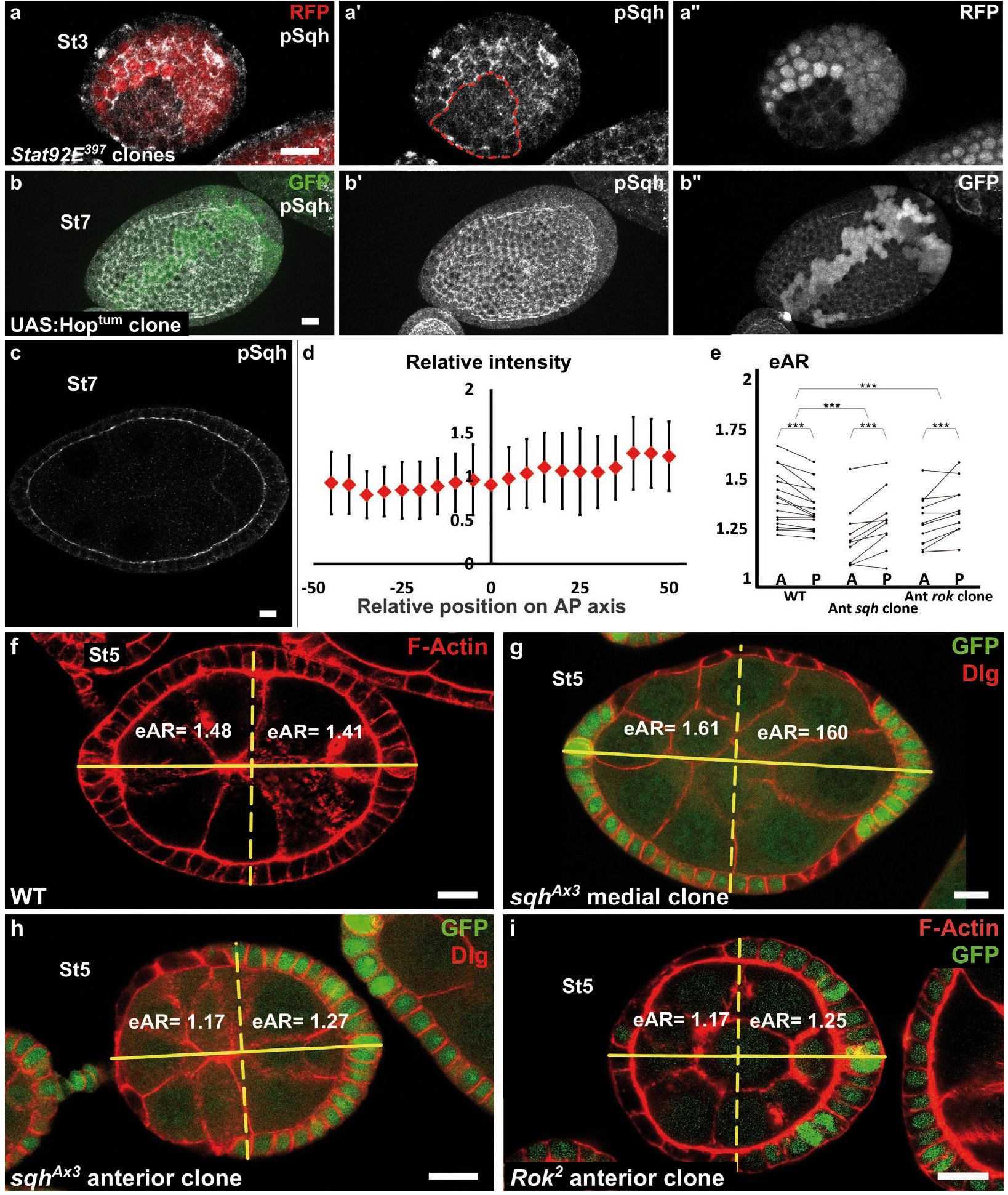
Myosin II is not controlled by JAK-STAT but required at the poles. a) Apical level of phosphorylated Sqh (pSqh, white and a’) is reduced in a mutant Stat92E clone (RFP-negative) in a stage 3 follicle (z-projection of the superior half of the follicle). b) Clonal overexpression of Hop^tum^ (green cells) on a stage 7 follicle is not sufficient to increase the expression of apical pSqh (white and b’) z-projection of the superior half of the follicle). c) pSqh staining in the middle plane of on a wild-type stage 7 follicle d) Quantification of the intensity of apical pSqh along the AP axis of stage 6-7 follicles. n = 5 follicles. Baseline value = mean apical pSqh per follicle. e) Quantification of the extrapolated aspect ratio (eAR) of stage 4 to 7 WT follicles or follicles with a *sqh*^A×3^ or *Rok*^2^ clone covering the anterior pole. Whereas, in WT follicles the anterior is significantly more curved than the posterior, the tendency is opposite with *sqh* and *Rok* clones. (p ***<0.001). f,g,h,i) representative images of f) WT g) mediolateral *sqh*^A×3^ clone h) anterior *sqh*^A×3^ clone, i) anterior *Rok*^2^ clone with the corresponding eARs.

If the gradient of apical pulses induces early elongation and explains MyoII involvement in this process, then MyoII function should be required at the poles. We generated mutant clones for a null allele of *sqh* to analyze where MyoII is required for elongation. As previously shown (Wang, & Riechmann 2007), such clones reach a limited size, probably explaining why it is rare to obtain a clone that covers poles, especially after stage 5. We focused on clones covering the anterior pole. To quantify the effect of mutant clones on semi-follicles, we measured each semi-follicle extrapolated Aspect Ratio (eAR), which means, the ratio of the corresponding full ellipse (see methods and Fig 5S1). For WT follicle, anterior eAR is equal or superior to posterior eAR, as the anterior pole is normally more pointed than the posterior (Fig 5f). Analysis of the eAR of the poles containing such mutant clones indicates that Myosin II loss of function specifically affects the elongation of this pole, compared to the opposite WT posterior poles (n=10) (Fig 5e,h). Moreover, we never observed clones in the mediolateral regions inducing elongation defects (n=35) (Fig 5g). Finally, we also performed similar experiment with *Rok null* mutant clones. Such clones have a weaker effect on cell morphology (Fig 5i and Wang, & Riechmann 2007), but still affect elongation when situated at the pole (Fig 5e,i). Thus, MyoII and Rok are required specifically at the poles to induce early elongation. These results strongly argue for the gradient of apical isotropic FC pulses as the force-generating mechanism that drives early elongation.

### Early elongation is associated with cell constriction and cell intercalation

Independently of the upstream events, we asked which cellular behavior was associated with early elongation. The simplest possibility would be that cells are stretched along the AP axis. However, cells are actually slightly elongated perpendicularly to the axis of elongation and this morphology did not change significantly over time, indicating that this parameter does not contribute to follicle elongation during early stages (Fig 6S1a,b). Tissue elongation can be also associated with oriented cell divisions. A movie of mitosis in the FE showed that this orientation is really variable through the different steps of mitosis (Fig 6S1c). We therefore quantified the orientation of cytokinesis figures, which did not highlight any bias towards the AP axis (Fig 6S1d). Finally, we asked whether early elongation could be associated with cell intercalation. Analysis of fluorescence video-microscopy images gave inconclusive results because such events are probably rare and slow and follicle rotation precludes their reproducible observation (Movie S13). We used therefore an indirect method. As from stage 6 follicle cells stop dividing and their number remains constant, we counted the number of cells in the longest line of the AP axis (i.e., the follicle plane that includes the polar cells). This number significantly increases between stage 6 and 8, showing that cells intercalate in this line (Fig 6a-d). This number was also correlated with the follicle AR (Fig 6e), indicating that follicle early elongation is associated with cell intercalation along the AP axis. Cell intercalation can be powered at a cellular scale by the polarized enrichment of Myosin II in the cells that rearrange their junctions (Bertet et al 2004). However, we have already shown that MyoII does not show such a pattern in FCs (Fig 3i,j). Alternatively, intercalation can be promoted at a tissue scale. For instance, apical cell constriction in the wing hinge induces cell intercalations in the pupal wing (Aigouy et al 2010). We observed that the cell apical surface is lower at the poles than in more equatorial cells, and that this difference increases during early elongation phase (Fig 6f,g,h). Such difference could be explained by cell shape changes or a differential cell growth. Cell height is significantly larger at the poles, indicating that the changes in apical surface are linked to cell morphology, as previously shown for instance during mesoderm invagination (Fig 6i) (He et al 2014). However, cells at the poles have a lower volume than in the mediolateral region at stage 7 (Fig 6S1e). This difference of volume is nonetheless proportionally weaker than the change in apical surface, suggesting the cell shape changes induce the reduction of volume rather than the opposite. Thus, early elongation is associated with a moderate cell constriction in the polar regions. *sqh* mutant FCs are stretched by the tension coming from germline growth, a defect opposite to cell constriction (Fig 5g,h)(Wang, & Riechmann 2007). Interestingly, FCs mutant for *Stat92E* are also flattened, with a larger surface and a lower height, compared to WT surrounding cells (Fig 6j-m). Moreover, the apical cell surface at the poles of stages 7-8 is increased by the loss of function of Upd (fig 6h). Hence, these results link JAK-STAT and the morphology of the follicle cells in a coherent manner with an involvement of apical pulses for the cell constriction observed at the poles.

**Figure 6:**
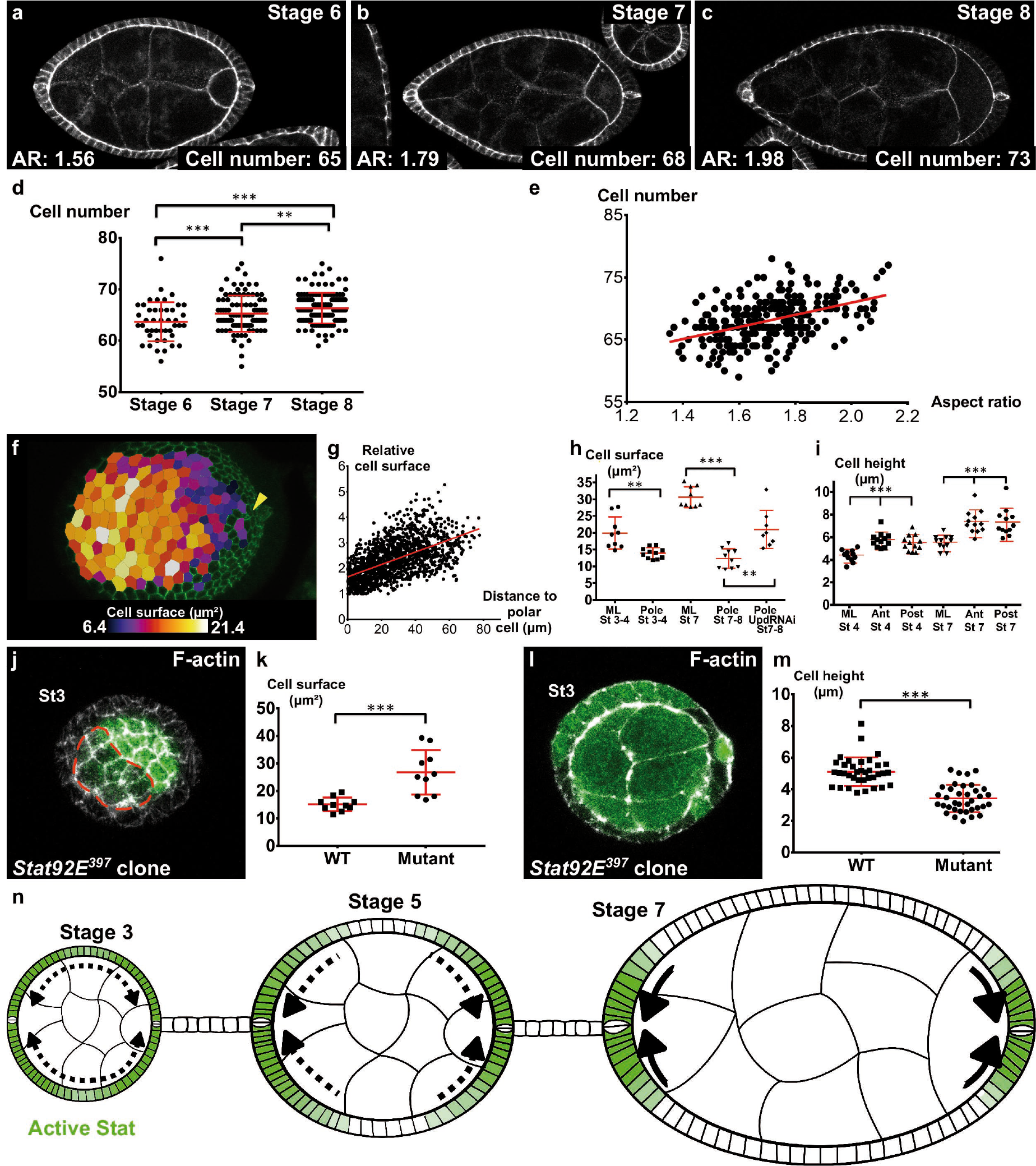
localized apical cell constriction and oriented cell intercalation occur during early elongation. a-e) Number of follicular cells in the plane of polar cells based on DE-Cad staining of stage 6 to 8 follicles depending on d) the stage and e) the aspect ratio of the follicle. a-c) AR and number of cells along the polar cell plane in representative stage (a) 6,(b) 7 and (c) 8 follicles stained for DE-Cad. f) Heat map of the cell apical surface of a representative stage 7 follicle imaged as on Fig 4g. Arrohead shos polar cells. g) Quantification of the relative apical cell surface (smallest cell = 1) in function of the distance from polar cells (n=10 stage 7 follicles, 1487 cells). h) apical cell surface and i) cell height depending on stage, position and genotype (ML: mediolateral). j) representative top view and l) section view of a *Stat92E*^397^ mutant clones at stage 3. Mutant cells have k) a larger apical surface and m) a loer cell height than wild-type cells. n) Schematic figure shoing the progressive restriction of JAK-STAT signaling (green) and of cell constriction to the follicle poles. p *< 0.05,**<0.01,***<0.001.

Thus, altogether these results indicate that two cell behaviors occur during the early phase of elongation: oriented cell intercalation towards the A-P axis and apical cell constriction at the poles.

## Discussion

The first main conclusion of this work is that follicle elongation can be subdivided in at least two main temporal and mechanistic phases: an early one (stage 3 to 7) that is independent of Fat2, rotation and ECM and F-actin basal polarization, and a second one (stage 8 to 14) that requires Fat2. This is reminiscent of germband extension where different elongation mechanisms have been described (Lye et al 2015; Collinet et al 2015; Rauzi et al 2010; Blankenship et al 2006; Sun et al 2017). In the case of the follicle, it is still not clear how overlapping and interconnected these different mechanisms are.

Fat2 has no role in early elongation. Nevertheless, Fat2 is required as early as the germarium for the correct planar polarization of the microtubule cytoskeleton and for follicle rotation, that takes place during the early elongation phase (Viktorinová, & Dahmann 2013, Chen et al 2016). The rotation reinforces the basal pcp of the F-actin during stages 4 to 6, and thus likely participates to the late phase in this way (Cetera et al 2014; Aurich, & Dahmann 2016). Rotation is also necessary for the ECM fibrils deposition, though, their specific role in elongation has not been clearly elucidated yet. Another mechanism participating to elongation is the ECM stiffness gradient (Crest *et al*, 2017). However, its contribution begins only at stage 7-8, in agreement with the fact that it depends on Fat2 and that *vkg* (ColIV) loss of function follicles elongate correctly up to stage 8, showing that the ECM is required only in the second elongation phase (Crest et al 2017; Haigo, & Bilder 2011). Thus, the setting up of the elements required for this second elongation phase fully overlaps with the first elongation phase, but these two phases are so far unrelated at the mechanistic level. Notably, the early elongation phase requires elements of the apical side of follicle cells, whereas the second phase involves the basal side. Mirroring our observations, a recent report nicely shows that the fly germband extension, which was thought to depend exclusively on the apical domain of the cells, also involves their basal domain (Sun et al 2017). Since both Fat2 and the gradient of BM stiffness are involved in the elongation at stage 8 and that apical pulses are still observed at this stage, it suggests that apical and basal domain contributions may slightly overlap. Moreover, both the gradients of apical pulses and of BM stiffness are under the control of JAK-STAT, indicating that this pathway has a pleiotropic effect on follicle elongation.

We have also shown that integrin and *Pak* contribute to early elongation in an indirect manner through their impact on the positioning, the differentiation or the survival of the polar cells. In this respect, *Pak* and *mys* mutants belong to a new phenotypic class that could also comprise the Laminin β1 subunit (LanB1) and the receptor-like tyrosine phosphatase Lar (Díaz de la Loza et al 2017; Frydman, & Spradling 2001). It is yet unknown how the A-P position of those cells is established and maintained. Interestingly, *Pak* mutants have also an altered germarium structure leading to abnormal follicle budding, suggesting that polar cell mispositioning might be linked to this primary defect (Vlachos et al 2015). However, it is worth noticing that *Pak* mutant follicles do not elongate at all, whereas they still have a cluster of polar cells. Thus, *Pak* might be also required for early elongation in a more direct manner than polar cell positioning, downstream or in parallel to JAK-STAT pathway, but independently of basal planar polarization.

We found that polar cells define the elongation axis of each follicle during early elongation by secreting the Upd morphogen and forming a gradient from each pole, which in turn induces apical pulses. The isotropic nature of these pulses does not provide an evident link with tissue elongation, unlike the oriented basal pulses going on in later stages (He et al 2010). Moreover, the absence of planar polarization of MyoII in apical, the driving force of early elongation, and the non-requirement for “basal pcp” strongly argues against a control of this elongation phase via a planar cell polarity working at a local scale. Rather, several strong arguments allow proposing that the early elongation relies on pulses working at a tissue scale (Fig 6n). First, the pulses are distributed in a gradient from the poles, suggesting that this distribution can orient the elongation in each hemisphere. Also, our data indicate that JAK-STAT does not directly regulate MyoII activity, and, thus, they likely work in parallel to control pulses. The convergence of requirement of JAK-STAT and myosin II for both pulses and early elongation argues for a causal link between these two processes. To date, JAK-STAT has no other known morphogenetic function before stage 8. Similarly, the only other known function of MyoII is linked to the rotation, which is not involved in early elongation, and MyoII is very concentrated at the apical cortex, emphasizing the role of this domain. Moreover, though present all around the follicle, MyoII is required for early elongation at the poles. Thus, the apical localization and the spatiotemporal requirement of MyoII are coherent with the apical pulses acting as the driving force for early elongation.

JAK-STAT has been already involved in the elongation of different tissues in flies and in vertebrates. For instance, Upd works as the elongation cue for the hindgut during fly embryogenesis, a process also associated with cell intercalation, though the underlying mechanism is unknown (Johansen et al 2003). Maybe more significantly, JAK-STAT is involved in the extension-convergence mechanism during zebrafish gastrulation (Yamashita et al 2002). Moreover, JAK-STAT also participates in other morphogenetic events, such as tissue folding in the fly gut and wing disc (Wells et al 2013). All these roles are potentially linked to a control of apical cell pulses. Since our results indicate that this control is not through MyoII activation, identifying the transcriptional targets of STAT explaining its impact on apical actomyosin will be relevant for many developmental contexts.

How the apical pulses precisely drive early elongation remains a question that will require further investigations. Nonetheless, we determined that early elongation is associated with apical cell constriction close to the poles and oriented cell intercalations. Cell constriction is likely a direct consequence of apical pulses, as it has been shown in many other contexts, because both myosin II and JAK-STAT loss of function affect pulse and induce an increase of the apical surface (Wang, & Riechmann 2007; Martin, & Goldstein 2014). Thus, as during tissue invagination, cell constriction may accentuate the curvature at the poles and thus promote elongation. Intercalation can be induced at a tissue scale by long range anisotropic tensions in the tissue, as exemplified by pupal wing development or mammalian limb bud ectoderm (Aigouy et al 2010; Lau et al 2015). In the wing, elongation is due to contraction of the hinge, which corresponds to an apical constriction of the cells. Here, the apical pulses could act in a similar way via the constriction, acting as a pulling force at each pole. Thus, intercalations may correspond to a passive response bringing plasticity to the tissue, hence stabilizing its elongation. Although the respective contribution of these two cell behaviors - apical constriction at the poles and cell intercalation along the AP axis - and their potential links remain to be determined, together they likely recapitulate at the cellular scale the elongation observed at the tissue scale. Importantly, such a mechanism does not require any planar cell polarization, in agreement with our observations. To our knowledge, vertebrate AP axis elongation, which relies on a gradient of randomly oriented cell migration, is the only other example of tissue elongation instructed by a signaling cue and independent of planar cell polarization (Bénazéraf et al 2010). Our work proposes an alternative mechanism explaining how a morphogen gradient can induce elongation solely through transcription activation, and without any requirement for a polarization of receiving cells. This simple mechanism may apply to other tissues and other morphogens.

## Methods

### Genetics

All the fly stocks with their origin and reference are described in supplementary file 1A.

The detailed genotypes, temperature and heat-shock conditions are given in supplementary file 1B.

### Immunostaining and imaging

Dissection and immunostaining were performed as described previously (Vachias et al 2014) with the following exceptions: ovaries were dissected in Supplemented Schneider, each ovarioles were separated before fixation to obtained undistorted follicles. Primary antibodies were against pMyoII (1/100, Cell Signaling #3675), DE-Cad (1/100, DHSB #DCAD2), Dlg (1/200 DHSB #4F3), FasIII (1/200, DHSB #7G10). Images were taken using a Leica SP5 or SP8 confocal microscope. Stage determination was done using unambiguous reference criteria, which are independent of follicle shape (Spradling, 1993).

For live imaging, ovaries were dissected as described previously (Prasad et al 2007) with the following exceptions: each ovariole was separated on a microscope slide in a drop of medium and transferred into a micro-well (Ibidi BioValey©) with a final insulin concentration of 20μg/ml. Samples were cultured for less than 2 hours before imaging with a Leica SP8 confocal using a resonant scanner. Follicles were incubated with Y-27632 (Sigma) (diluted in PBS to 250 μM) for 10 to 30 minutes before image acquisition. To image the poles, glass beads were added in the well to form a monolayer (Sigma-Aldrich, G4649 for stage 6 to 8 or G1145 for earlier stages). Ovarioles were added on top of the beads and follicles falling vertically between the beads were imaged.

Cell pulse analysis was performed using the Imaris software and a MATLAB homemade script to segment and measure the cell surface on maximum intensity projections of 40 stacks taken every 15 seconds. The intensity of one cell pulsation corresponds to: (maximum surface of the cell - min surface)/(mean surface). The isotropy of one cell pulse is measured by dividing the AP and ML bounding box (best fit rectangle) axis length at cell maximal area by the AP and MP bounding box axis length respectively at cell’s minimal area. For each follicle, at least 10 cells were analyzed. For visualization (images presented in Fig 4a,c,d and the attached movies), the original files were deconvolved, but all the analyses were done using the raw files.

The Fiji software was used to measure the length of the long and short axis of each follicle on the transmitted light channel, and then to determine the aspect ratio in WT and mutant follicles. Cells in the longest line of the AP axis were counted manually using Fiji on the DNA and DE-Cadherin channels. Bazooka-GFP and MyosinII-mCherry enrichment were analyzed using the Packing Analyser software (Aigouy et al 2010). Cells were semi-automatically segmented based on the Baz-GFP channel that was used as common pattern to calculate the intensity of each bond for both channels. Fiji was used to measure the intensity of the pSqh signal and 10XStatGFP signal. A 15-pixel wide line was drawn using the freehand tool, either within the cells (10X StatGFP), or at the apical level of the cells (pSqh), from the anterior to the posterior of cross section images of follicles.

The extrapolated aspect ratio (eAR) was estimated for each pole by measuring the width of the follicle at 25% of its total length: for any given ellipse, this value corresponds to 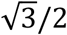 times its total width. Therefore, this measure allows, for each pole, to extrapolate a width and an aspect ratio. Based on Dlg staining, follicles with gaps in the epithelium were excluded. To measure cell elongation, images of DE-Cadherin-GFP expressing follicles were semi-automatically segmented using the Packing Analyser software and for each follicle the elongation tensor was calculated. The elongation tensor was defined by the mean elongation of all the segmented cells (elongation magnitude) and the mean orientation.

The rose diagrams were generated with Packing Analyser; each bin represents a 10° range and the bin size is proportional to the number of acquired data. Cell volume was obtained by the multiplication of the mean surface and the mean height of the cells.

Figures were assembled using ScientiFig (Aigouy, & Mirouse 2013).

### Statistical analysis

For all experiments, sample size is indicated in the figure legends or in supplementary file 1B. No statistical method was used to predetermine sample size. Results were obtained from at least two independent experiments, and for each experiment multiple females were dissected. No randomization or blinding were performed. For each experimental condition variance was low. Matlab software has been used to make analysis of covariance to determine the elongation coefficient and a multiple pairwise comparison test has been run to determine the p-value between different conditions (aoctool and *multicompare*, Statistic and Machine Learning Toolbox). The normality of the samples has been calculated using a D’Agostino & Pearson normality test. Unpaired t-test has been used to compared samples having a normal distribution. Unpaired Mann-Whitney test has been used to compared samples having non normal distribution. For comparison of eAR of anterior and posterior poles, a two-way ANOVA test with repeated measures was conducted on both poles and for two genotypes. The post-hoc analysis (two pair-wise Bonferroni tests) was performed. When shown, error bars represent s.d. For all figures p *< 0.01, **<0.005, ***<0.001.

## Acknowledgements

We thank R Basto, M Crozatier, C Dahmann, M Grammont, D Harrison, A-M Pret and E Wieschaus for fly stocks or reagents. This work was funded by the ATIP-Avenir program, Association pour la Recherche contre le Cancer (ARC) and the Auvergne Region. We also thank the confocal imaging facility of Clermont-Ferrand (ICCF), and team members for comments on the manuscript.

### Competing financial interests

The authors declare no competing financial interests.

---

**Figure 1 – Figure S1:**
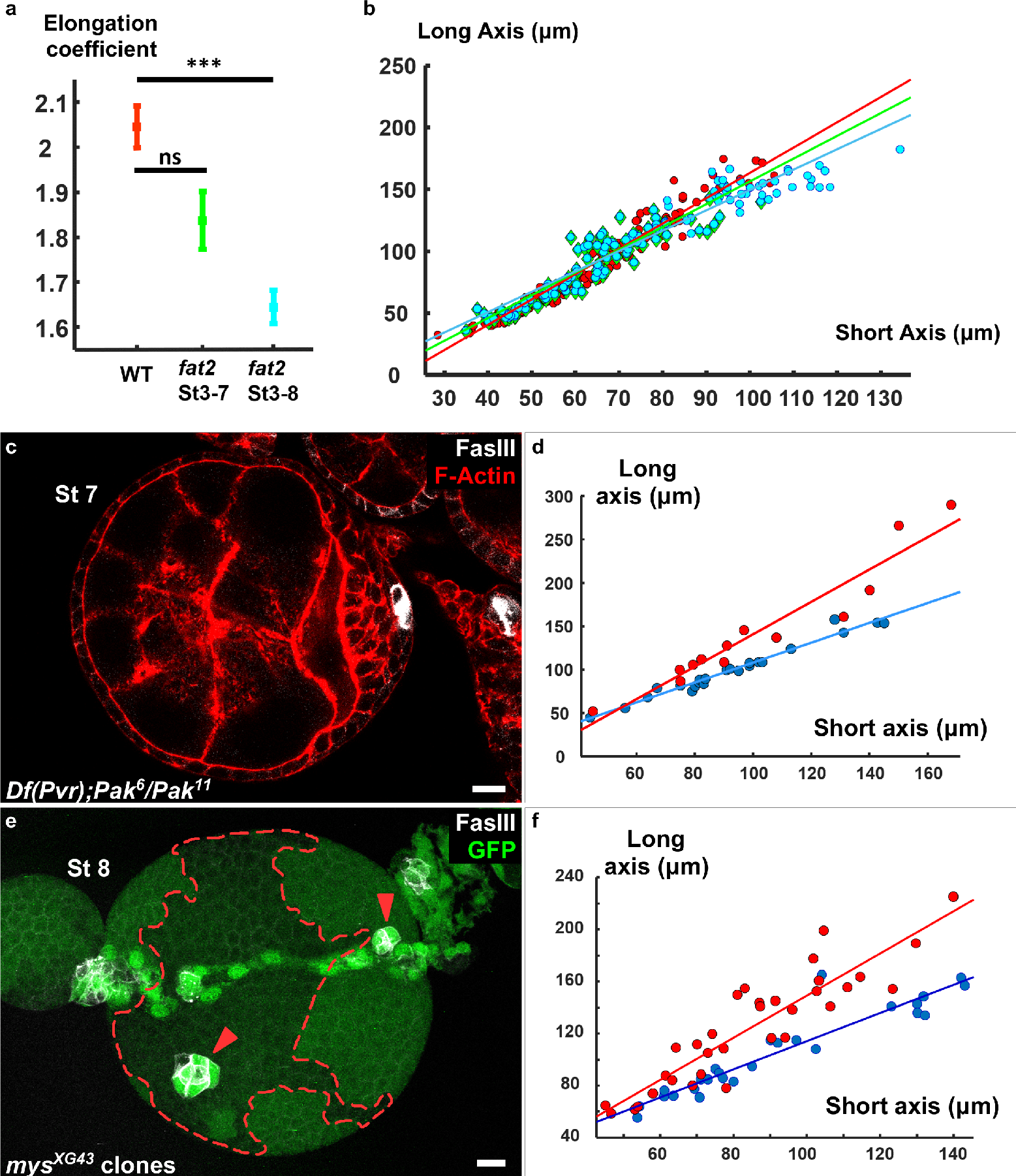
a) Elongation coefficients corresponding to the slope of the regression lines of the plots in b). b,d,f) plot of the long axis as function of a short axis of b) *fat2* mutant follicles (red = WT, green = *fat2* st3-7,blue *fat2* st 3-8) d) *Pak*^6^/*Pak*^11^,*Df*(*Pvr*)/+ follicles and f) follicles containing mutant clones for mys, affecting (blue) or not (red) polar cell positioning. Corresponding regression lines are represented. c) *Pak*^6^/*Pak*^11^,*Df*(*Pvr*)/+ follicle with a single cluster of polar cells. e) z-projection of a follicle with a *mys* mutant clone and two misplaced polar cell clusters (red arrowheads). The green signal in the mutant clone comes from germline signal due to the z-projection.

**Figure 2 – Figure S1:**
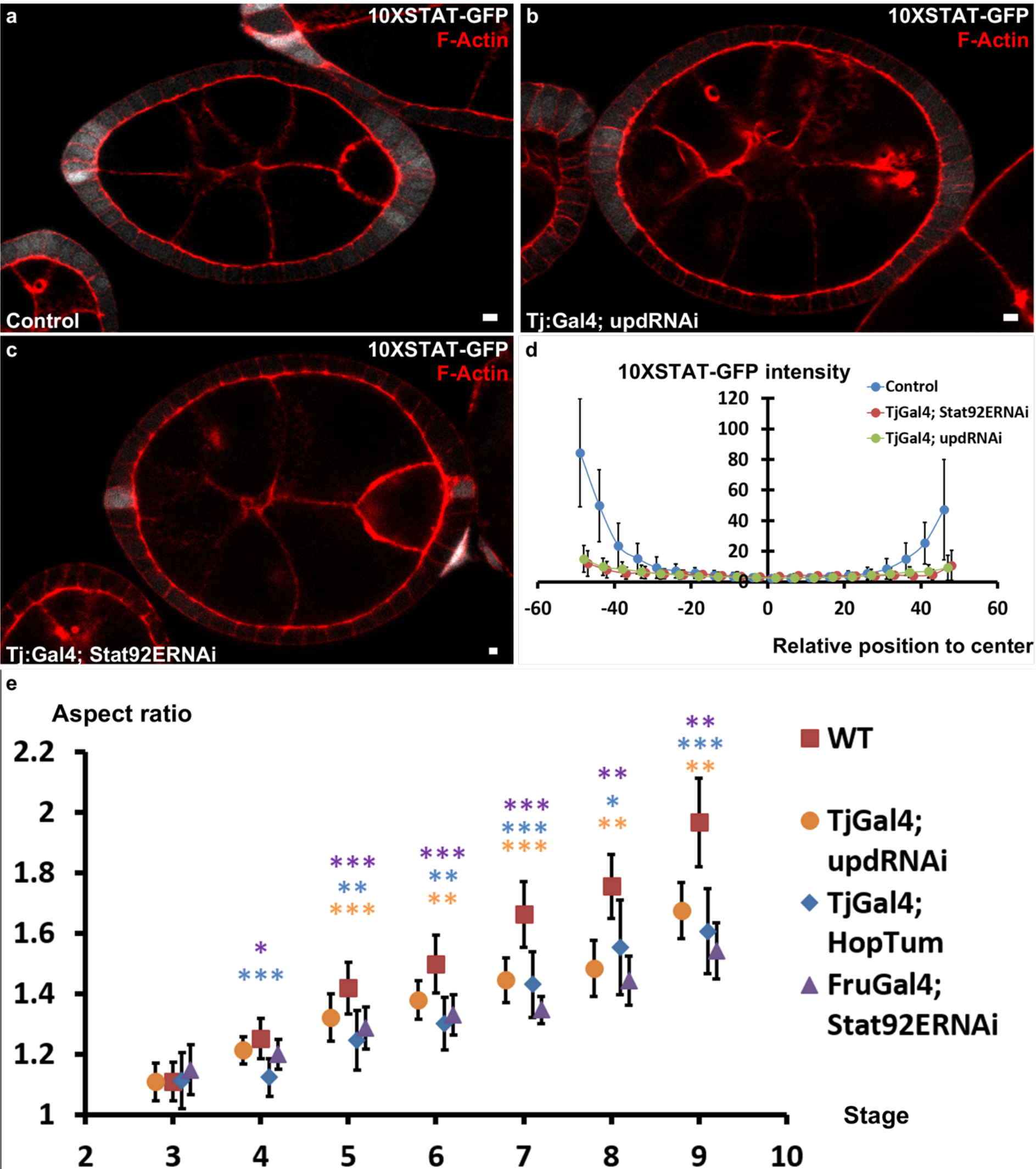
a to c) Representative follicles expressing 10XStatGFP in a) WT b) *upd RNAi* or Stat92E RNAi background. d) Quantification of the Stat activity gradient for the indicated genotypes at stage7 using the 10XSTATGFP reporter. e) Quantification of the aspect ratio (AR) in WT and the different JAK-STAT mutants (loss and gain of function) during early and intermediate stages of elongation (p * 0.05,**<0.01,***<0.001).

**Figure 3 – Figure S1:**
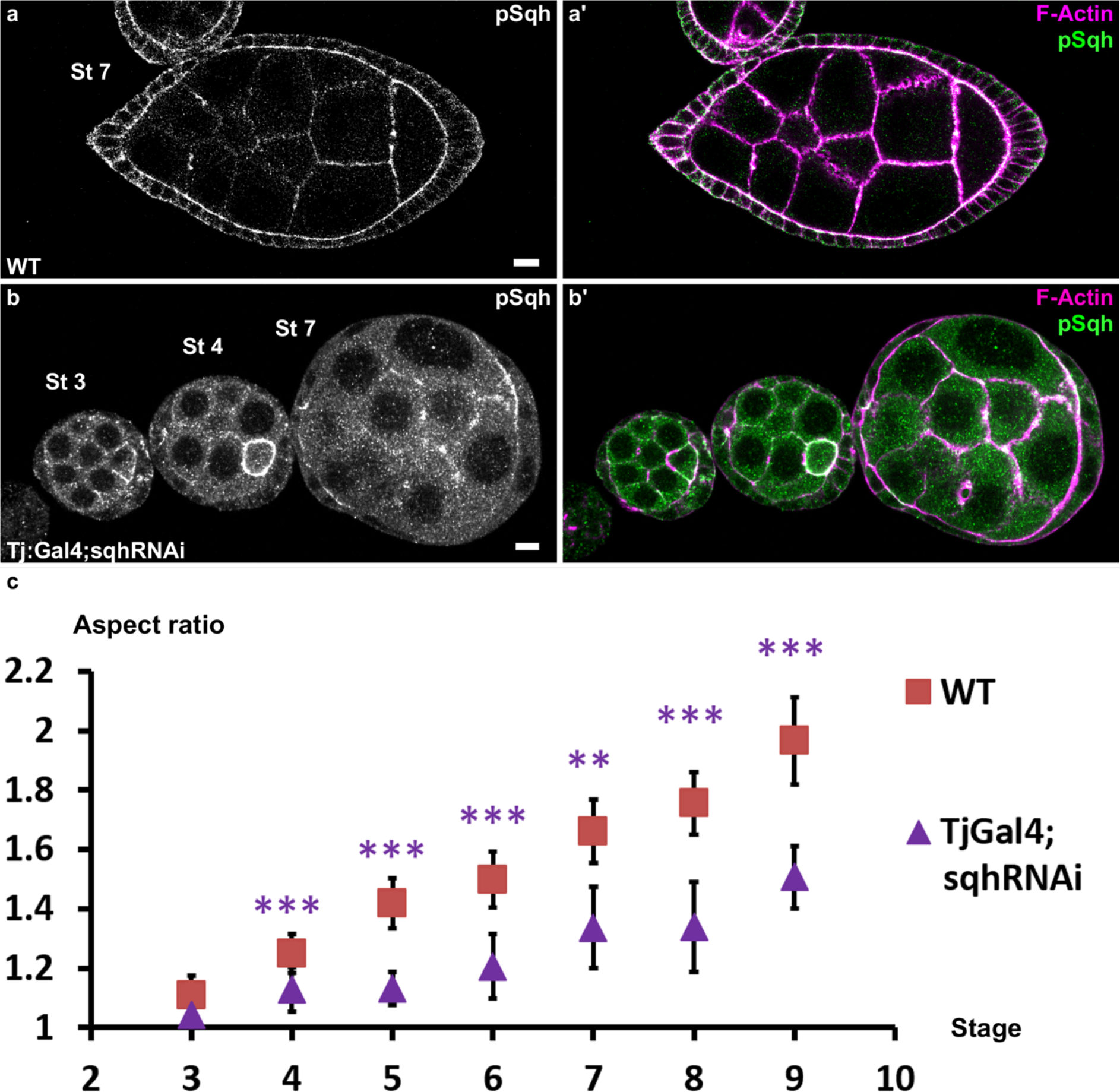
Follicles stained for pSqh (white in a,b, green in a’,b’) and F-actin (pink in a‘b’). a) WT follicle b) SqhRNAi in follicle cells driven with Tj:Gal4. The signal is mainly apical in WT follicle cells and is strongly reduced in the knock-down. c) AR quantification in WT or sqh knock-down follicles during the early elongation phase. (p ***<0.001)

**Figure 4 – Figure S1:**
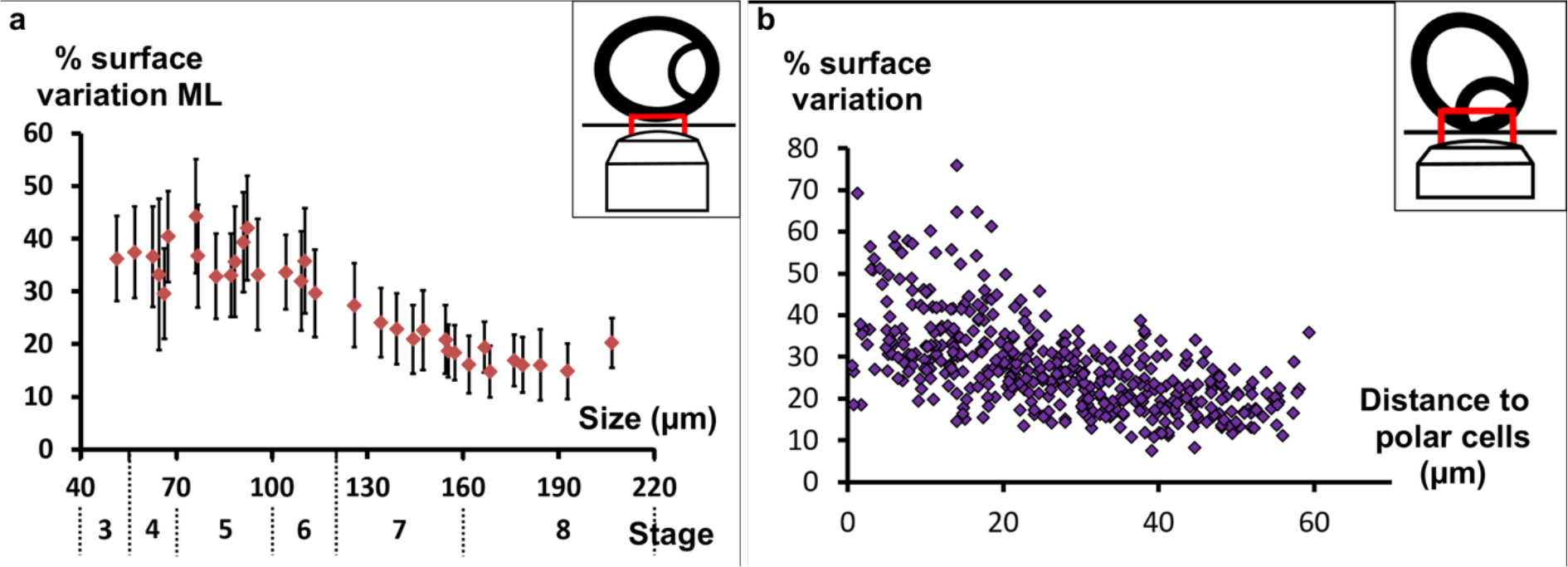
a) Mean percentage of apical surface variation in the mediolateral region relative to the follicle length (corresponding stages are also indicated). b) Pulsing activity of individual cells from five stage 7 follicles (n=441 cells) relative to their distance from the polar cells (tilted view from the pole).

**Figure 5 – Figure S1:**
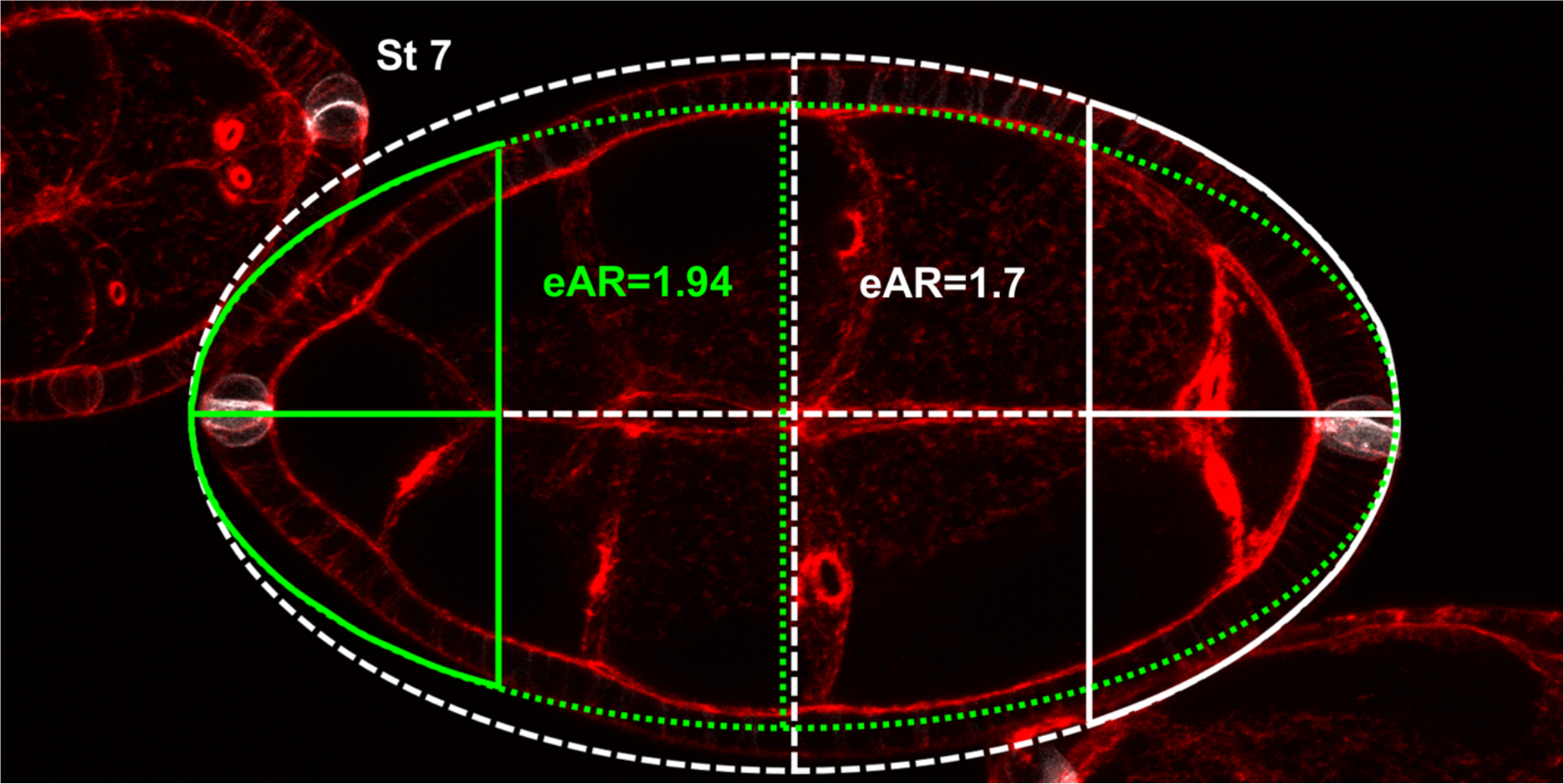
illustration of extrapolated Aspect Ratio (eAR) calculation based on width measure of a pole at 25% of AP axis length.

**Figure 6 – Figure S1:**
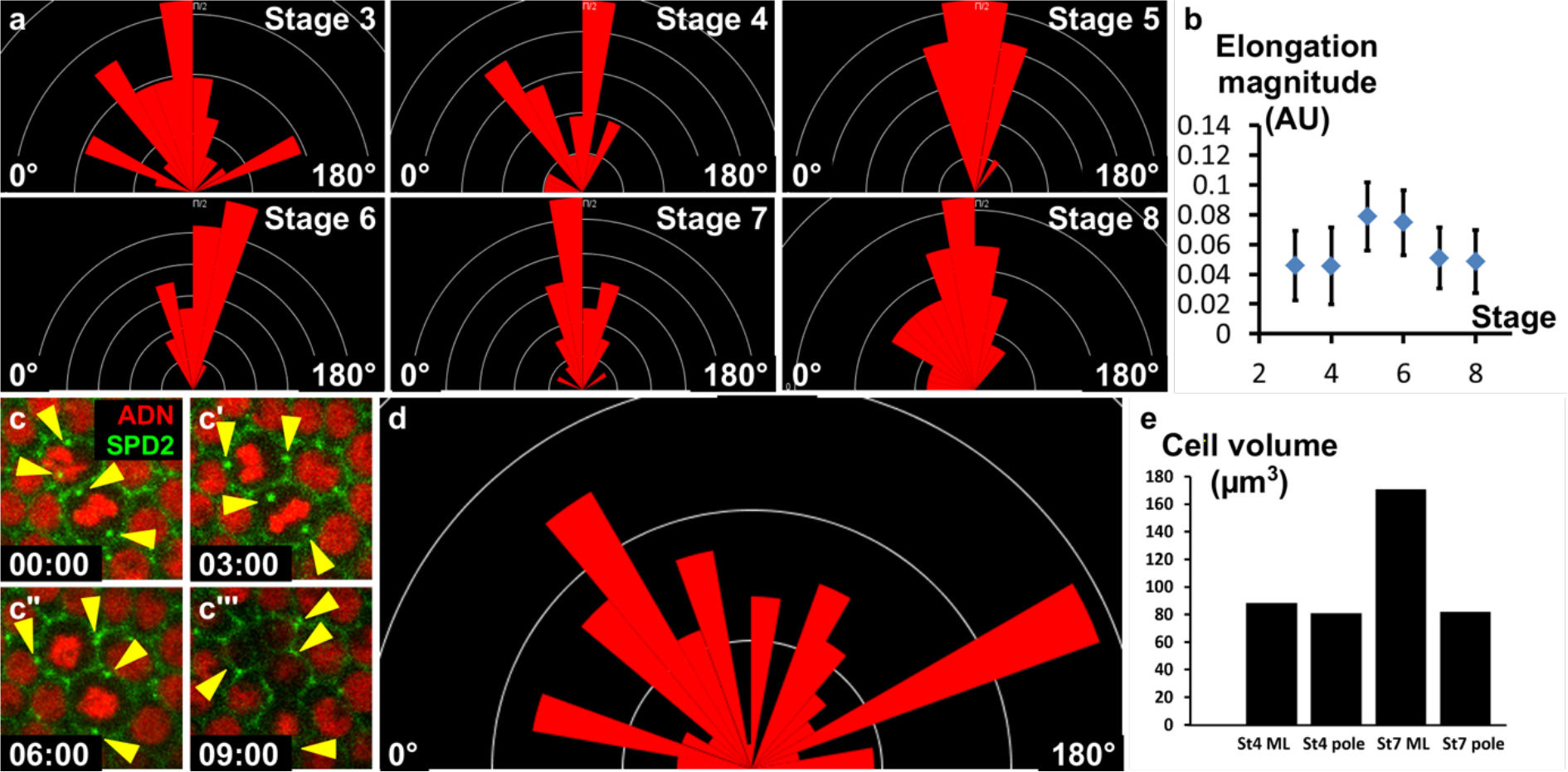
a) Orientation of cells with elongated shape in the mediolateral part of the follicle at the indicated stages (the X axis corresponds to the AP axis). b) Magnitude of the elongated cell shape according to the stage. Values correspond to an aspect ratio between 1.1 and 1.2. c) Fluorescence video-microscopy images of dividing wild type follicular cells that express H2A-RFP and SPD2-GFP to mark the centrosomes (yellow arrowheads). d) Orientation of cytokinetic figures (the X axis corresponds to AP axis). e) Calculation of FC volume depending on the position (pole or mediolateral (ML)) and the stage. Volume is higher in st7 ML due to the mitosis/endoreplication switch occurring at stage 6. However, cells at the poles maintains a lower volume.

## Movie description

### Movie S1

Full z-stack of a follicle with a *mys* mutant clone that affects polar cells

### Movie S2

Full z-stack of a follicle with a *mys* mutant clone that does not affect polar cells

### Movie S3

Stage 3 follicle expressing Sqh-GFP. The pool of apical myosinII is very dynamic.

### Movie S4

Zoom in on a cell of a stage 3 follicle expressing Baz-GFP and Sqh-mCherry. Apical MyosinII enrichment occurs at the same time as the apical cell domain contracts.

### Movie S5

Stage 3 follicle expressing Baz-GFP. Cells in the mediolateral part undergo apical pulsations.

### Movie S6

Stage 7 follicle expressing Baz-GFP. The apical surface variation is strongly reduced on the mediolateral part compared with stage 3 follicles.

### Movie S7

Stage 7 follicle expressing Baz-GFP observed from the pole. Polar cells are indicated on the corresponding Figure 5d (red arrowheads). The pulse intensity remains high in these cells compared with movie S4. The rotation is visible and occurs around the polar cells.

### Movie S8

Stage 3 follicle expressing Baz-GFP observed from the pole. Polar cells are indicated on the corresponding Figure 5b (red arrowhead).

### Movie S9

Stage 3 *upd* knock-down follicle expressing Baz-GFP. The intensity of the pulse is reduced compared with a WT stage 3 follicle (movie S3).

### Movie S10

Stage 7 *upd* knock-down follicle expressing Baz-GFP and observed from the pole. The intensity of the pulse is reduced compared with a WT stage 7 follicle (movie S6).

### Movie S11

Stage 7 follicle expressing ectopically Baz-mCherry. The intensity of the pulse is low.

### Movie S12

Stage 7 follicle expressing ectopically Baz-mCherry and Hop^tum^. Activation of the JAK-STAT pathway is sufficient to increase the pulsing.

### Movie S13

Movie representing a stage 7 DE-Cad-GFP follicle imaged during two hours. One raw of cell is tracked (red line). No intercalation occurs during this period.

**Supplementary table 1A.**
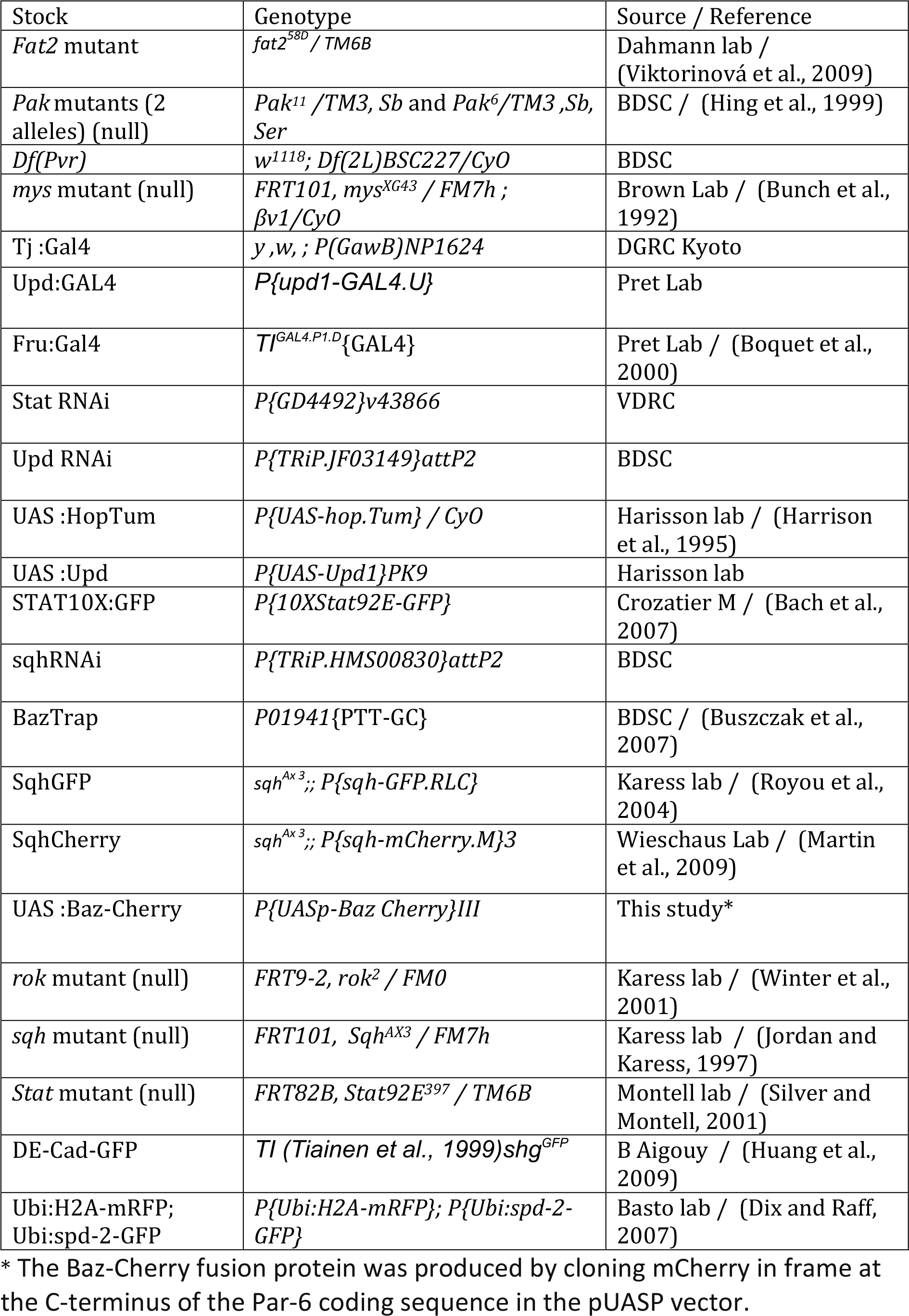
stock list and source

**Supplementary table 1B.**
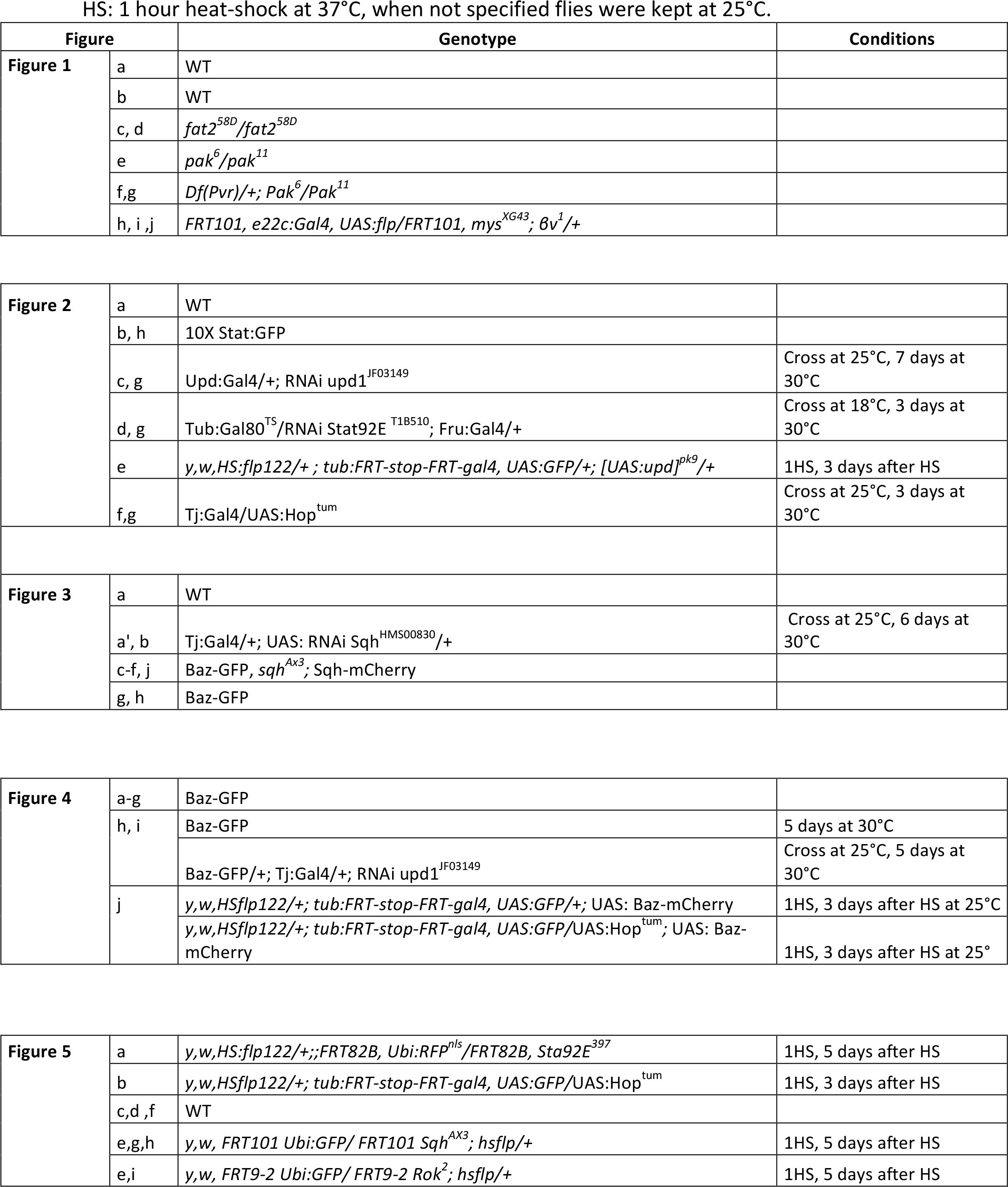
detailed genotypes and specific conditions

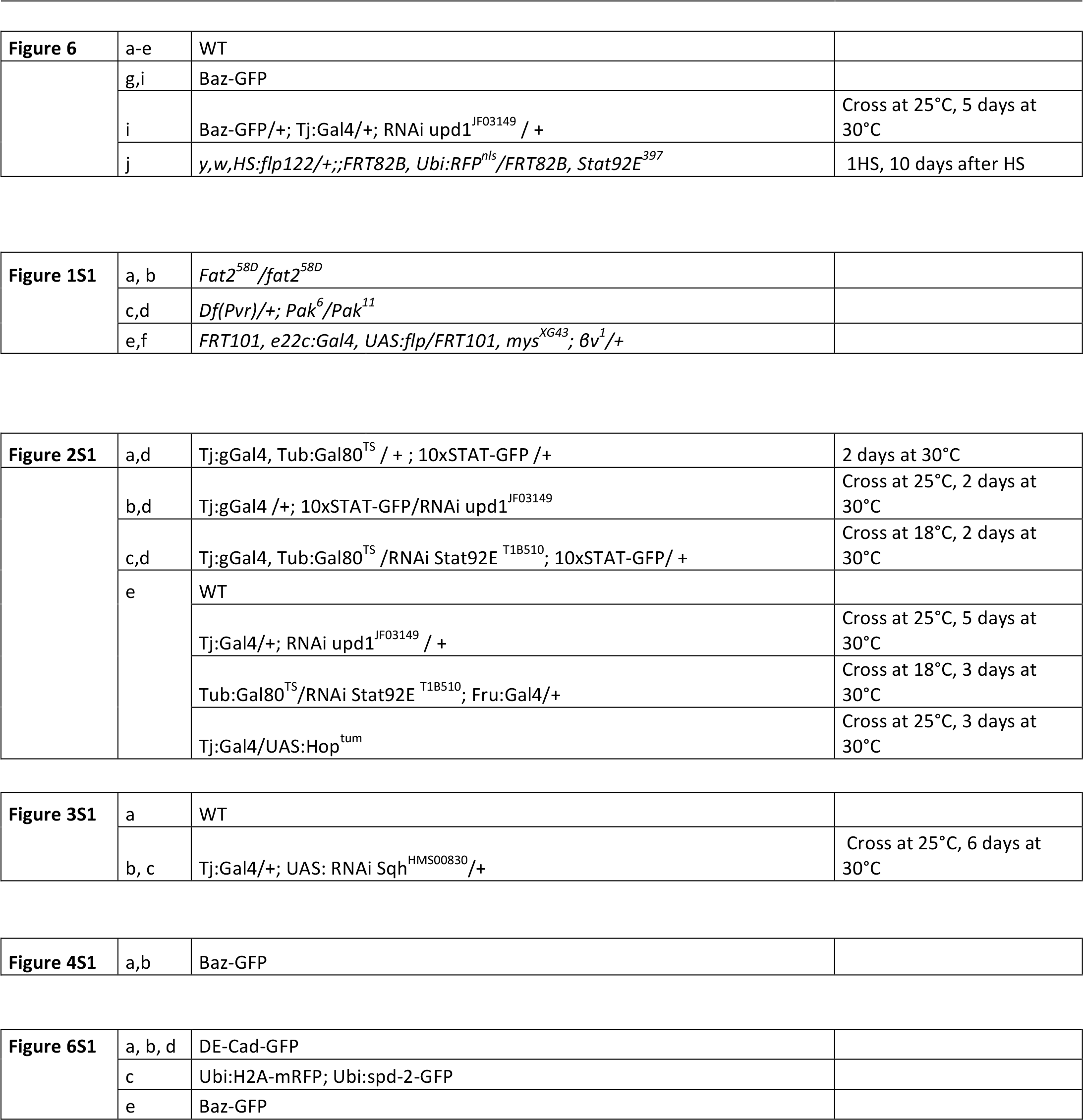

**Supplementary table 1C.**
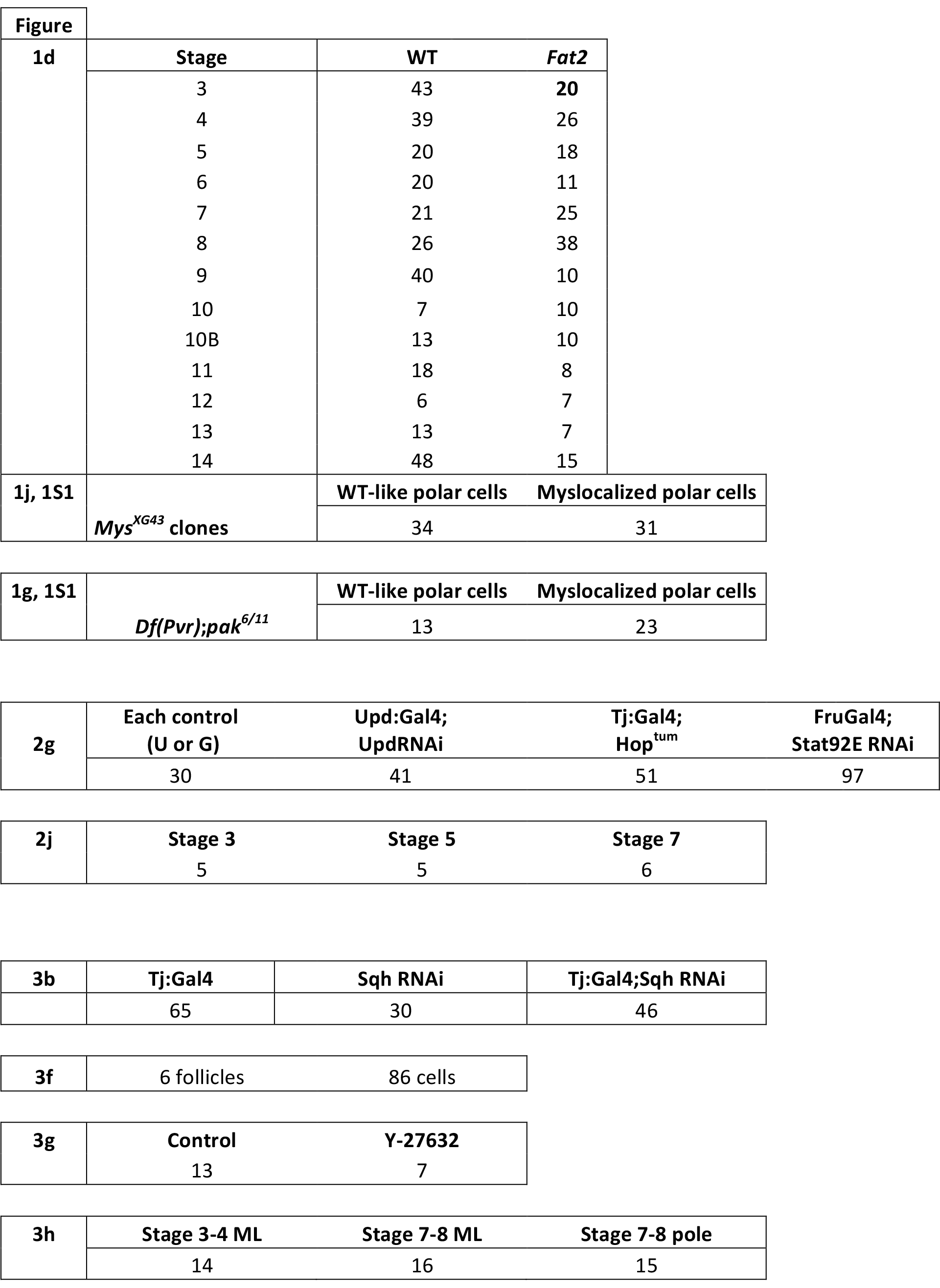
detailed sample size

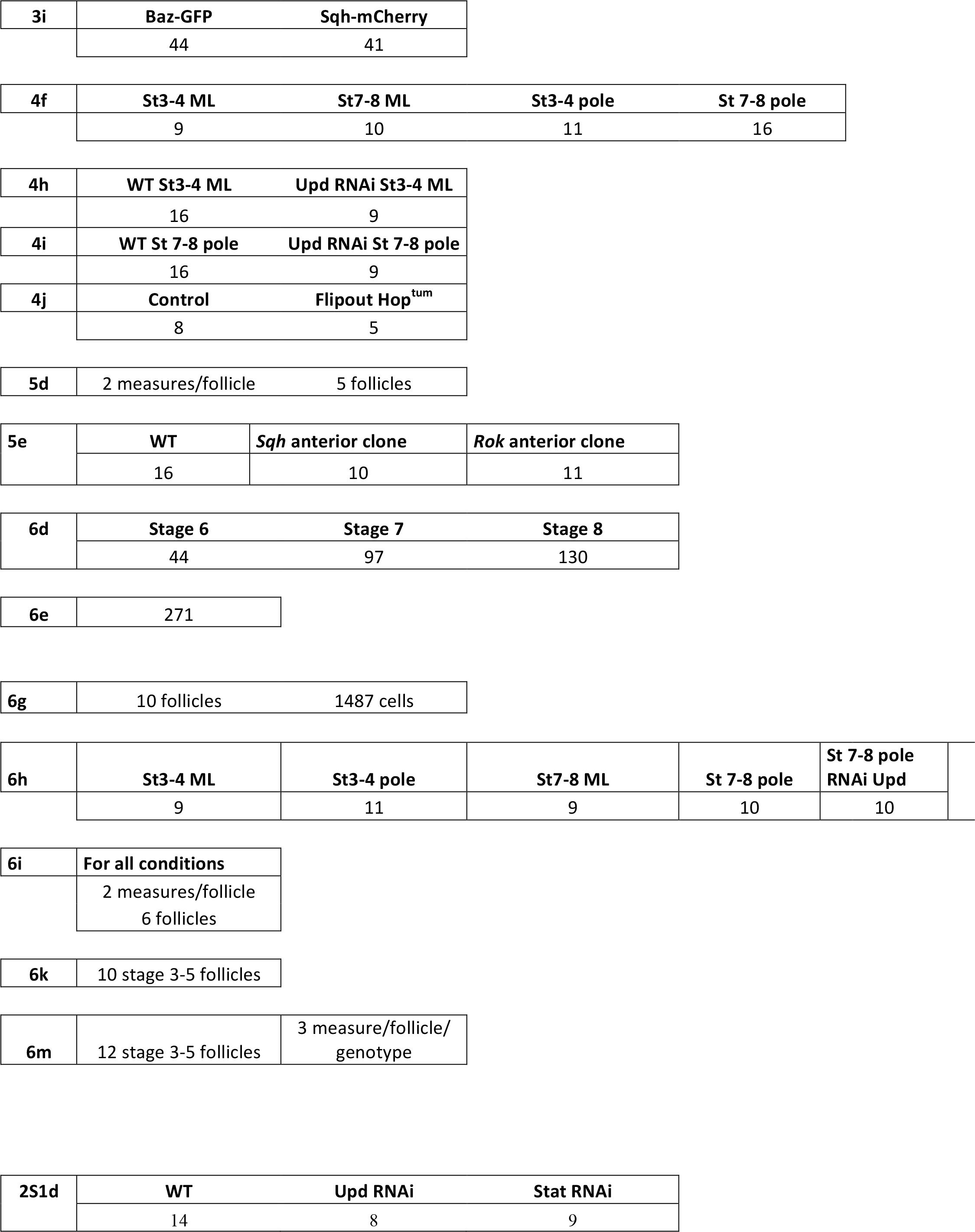

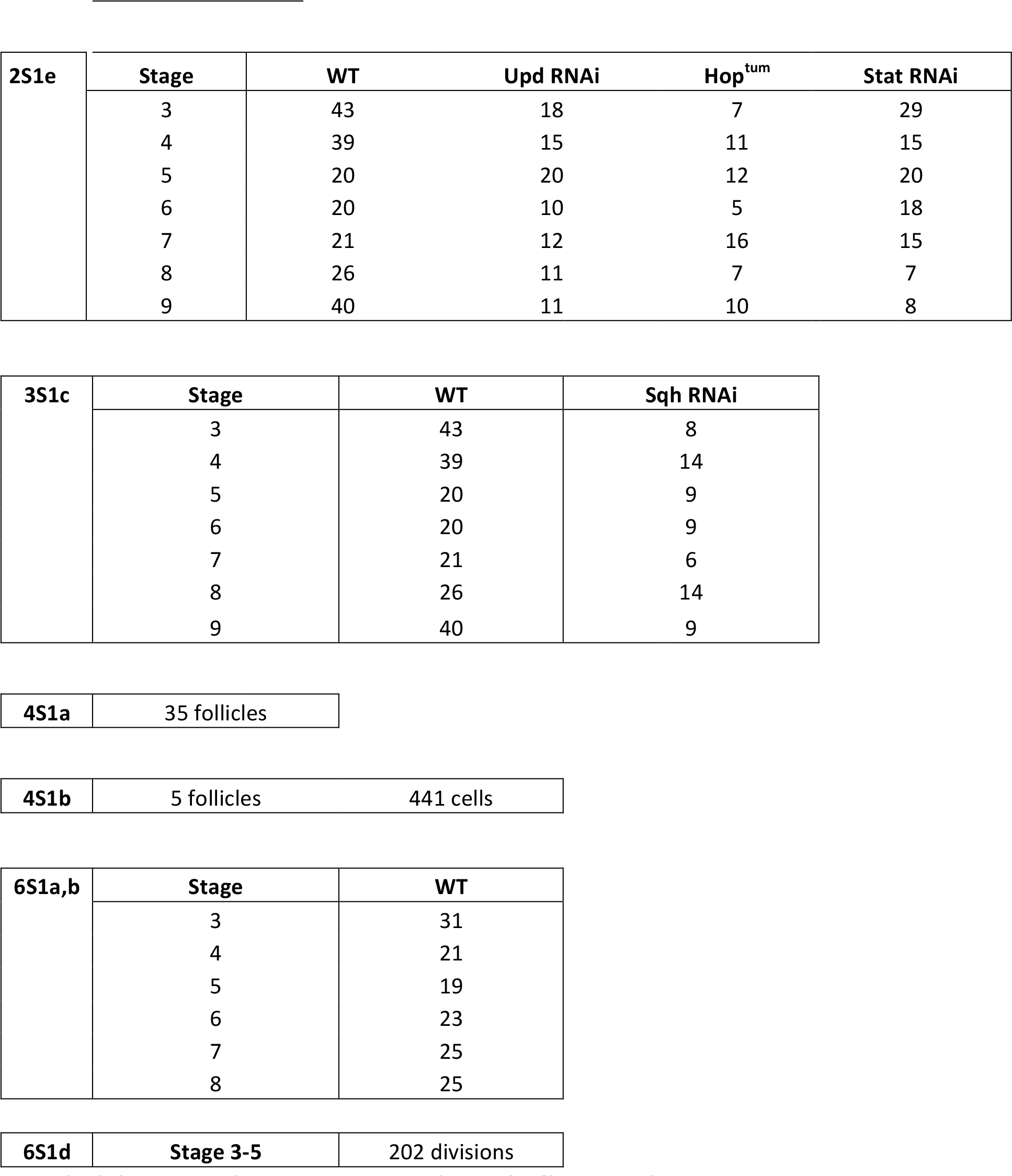

